# Evaluating the reliability of tools for mRNA annotation and IRES studies

**DOI:** 10.64898/2026.03.29.707813

**Authors:** Gemma E. May, Christina Akirtava, Joel McManus

**Affiliations:** Department of Biological Sciences, Carnegie Mellon University, Pittsburgh PA 15213; Department of Human Genetics, University of Utah School of Medicine, Salt Lake City, 84132; Computational Biology Department, Carnegie Mellon University, Pittsburgh PA 15213

## Abstract

Since the discovery of viral Internal Ribosome Entry Sites (IRESes), researchers have sought similar elements in mammalian genes, termed “cellular IRESes”. However, the plasmids used to measure cellular IRES activity are vulnerable to false-positives due to promoter activity in candidate IRESes. Orthogonal methods are needed to validate putative IRESes while carefully avoiding known artifacts. Recently, Koch et al. proposed approaches for studying IRESes, primarily circular RNA-generating plasmids, and for validating mRNA transcripts using smFISH and qRT-PCR. Here, we demonstrate confounding variables and artifacts in each approach that can lead to unwarranted conclusions about potential cellular IRES activity. We show the back-splicing circRNA plasmid creates linear mRNA artifacts associated with false-positive IRES signals. Using orthogonal assays validated with viral IRESes, we find putative cellular IRESes reported using the back-splicing plasmid have no IRES activity. Furthermore, we demonstrate that smFISH and qRT-PCR can misidentify nuclear ncRNAs as mRNAs and we validate a single molecule sequencing assay for identifying mRNA 5’ ends. Our work establishes reliable methods for robust transcript annotation and IRES studies that avoid documented artifacts.

## Introduction

Eukaryotic translation initiation requires recruitment of ribosomes to start codons. In most cases, 40S subunits are recruited to mRNA 5’ ends and scan directionally until a start codon is identified (Hinnebusch *et al*, 2016; Pelletier & Sonenberg, 2019). Ribosomes can also be recruited directly to start codons via Internal Ribosome Entry Sites (IRESes). First identified in poliovirus (Pelletier & Sonenberg, 1988; Jang *et al,* 1988), many viral IRESes have been thoroughly studied and validated. Because many gene expression processes first discovered in viruses were later found in human genes (Enquist, 2009), researchers have sought to find active IRESes in mammalian genes, referred to as “cellular IRESes”. Indeed, many studies reported cellular IRES activity using a variety of plasmid reporter strategies. These plasmids place candidate IRES sequences between two reporter genes (bicistronic) or in circular RNAs (circRNAs) generated by back-splicing, such that translation initiation by cap-dependent scanning should not occur. However, splice sites and transcriptional promoters in candidate IRESes can lead to false positives in *both* plasmid reporter systems (Thompson, 2012; Terenin *et al*, 2017; Chu *et al*, 2021; Ho-Xuan *et al*, 2020; Dodbele *et al*, 2021; Jiang *et al*, 2021; Unti *et al*, 2024). To avoid plasmid false-positives, direct transfection of mRNA constructs has become a gold-standard for IRES studies (Terenin *et al*, 2017; Loughran *et al*, 2025). Alternatively, the “Tornado” plasmid recently emerged as a reliable circRNA system that minimizes the promoter-driven artifacts generated by back-splicing circRNA plasmids (Unti *et al*, 2024).

The false-positive IRES problem is exacerbated by 5’ UTR annotation errors and the complexity of mammalian transcription and splicing (Akirtava *et al*, 2022). For example, a putative cellular IRES was reported in a long 5’ UTR of the mouse *Hoxa9* gene, based primarily on its activity in a plasmid bicistronic reporter assay (Xue *et al*, 2015). However, two studies found that the long *Hoxa9* 5’ UTR was misannotated and showed that the putative IRES was simply the *Hoxa9* promoter (Ivanov *et al*, 2022; Akirtava *et al*, 2022; Fig. EV1), suggesting the apparent IRES activities previously reported from this region resulted from promoter-driven artifacts. These studies also found trace amounts of RNA upstream of the *Hoxa9* promoter in public RNA-seq data and in calibrated qPCR experiments. It was proposed that these upstream RNAs were excised introns from known fusion transcripts between *Hoxa9* and *Hoxa10* (Mainguy *et al*, 2007) and primary precursor transcripts for the miRNA mir196b that splice into *Hoxa9* exons (pri-miR196b, Fig. EV1; Akirtava et al., 2022;(Rawat *et al*, 2020)). The presence of overlapping coding, fusion, and non-coding transcripts, some of which include the *Hoxa9* promoter region, likely caused the initial misannotation of the *Hoxa9* 5’ UTR. This complexity also complicates the interpretation of experiments using probes and primers targeting *Hoxa9*, as these could also detect and amplify shared sequences in pri-miR196b, excised introns, and fusion transcripts. Modern technologies that sequence full-length mRNA – from m7G-capped 5’ ends to polyadenylated 3’ ends – may be more reliable for mRNA annotation.

Recently, “A versatile toolbox for determining IRES activity in cells and embryonic tissues” (Koch *et al*, 2025; henceforth referred to as “the toolbox study”) described attempts to find reliable approaches to assay putative IRESes and identify bona fide 5’ UTRs. The authors promote back-splicing “circRNA” plasmids and qRT-PCR amplification across sucrose gradient fractions to assess IRES-driven translation. The toolbox study claimed its back-splicing plasmid separates IRES and promoter activity such that the presence of promoters in candidate IRESes is irrelevant. Finally, the study claimed that third generation full-length RNA sequencing does not reliably detect mRNA transcript 5’ ends and instead advocate for smFISH and qRT-PCR for 5’ UTR annotation.

Here, we identify confounding variables and artifacts in each of the systems described by the toolbox study that cause errors in mRNA transcript annotation and false-positive IRES activities. We show that the ZKSCAN1 back-splicing circRNA system is confounded by abundant *trans*-spliced linear artifacts that arise from transcriptional promoters in candidate IRESes from the toolbox study These *trans*-spliced linear artifacts allow for cap-dependent translation. The presence of these artifacts invalidates claims that promoter activities are irrelevant to this back-splicing circRNA plasmid. Importantly, using two orthogonal assays that we extensively validated with positive and negative controls, we show that three reported cellular IRESes from the toolbox study have no IRES activity. We also show that non-translating RNAs can contaminate polysome fractions in sucrose gradients, creating additional false-positive signals. Finally, thorough analyses of mouse embryonic PacBio Iso-Seq and smFISH data indicate the putative “IRES” isoform of *Hoxa9*, which includes the bona fide *Hoxa9* promoter, is not expressed as an mRNA in mouse embryonic tissues. Instead, the *Hoxa9* promoter region is transcribed as part of a pri-mir196b lncRNA and is detectable by smFISH only in embryo cell nuclei, not in the cytoplasm.

## Results

### The toolbox study’s back-splicing plasmid produces abundant *trans*-spliced linear GFP artifacts

A central claim in the toolbox study is that a back-splicing circRNA plasmid system (Chen *et al*, 2021) is a reliable IRES assay. This plasmid generates circRNAs by splicing a downstream 5’ splice site to an upstream 3’ splice site, driven by base pairing of complementary Alu elements in introns from the ZKSCAN1 gene (Fig. 1A). However, previous studies found similar back-splicing plasmids also make linear monocistronic transcripts that are the major source of reporter protein translation (Jiang *et al*, 2021; Chu *et al*, 2021; Ho-Xuan *et al*, 2020). Promoter activity in candidate IRES sequences drives the production of these monocistronic mRNA artifacts via *trans*-splicing (Fig. 1B). This *trans*-splicing can also be driven by Alu element pairing (in *trans* rather than in *cis*). Indeed, the creator of the ZKSCAN1 back-splicing plasmid system wrote:

**Figure 1.**
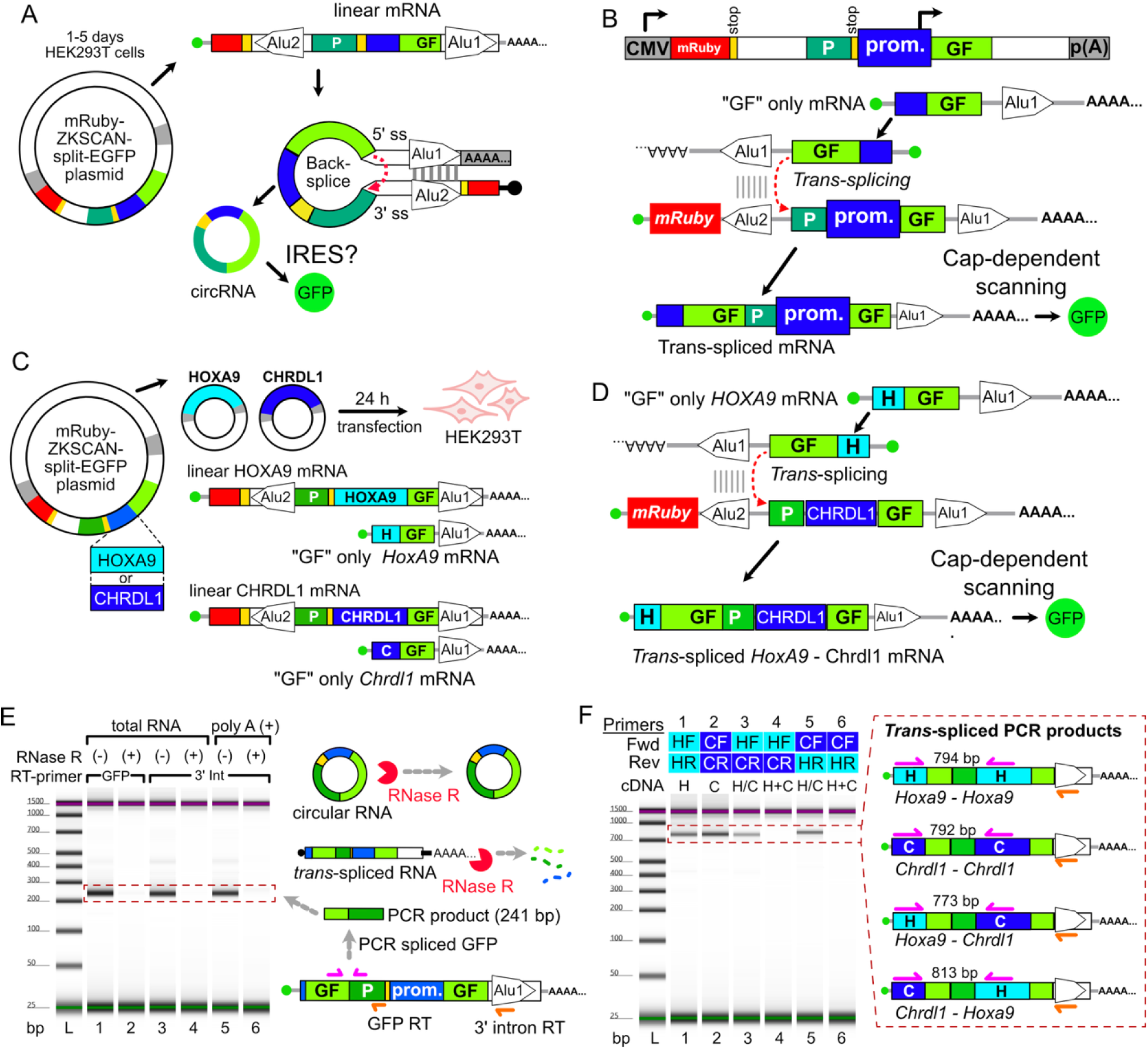
The back-splicing circRNA plasmid from the toolbox study produces linear trans-spliced artifacts. **(A)** Diagram of the ZKSCAN1 plasmid designed to generate circRNA by back-splicing. **(B)** Previous studies found ZKSCAN1 circRNA plasmids generated capped linear transcripts via trans-splicing that can express reporter proteins by cap-dependent translation. **(C)** Experimental design to test for trans-spliced artifacts. The Koch et al. reporter plasmids containing the Hoxa9 and Chrdl1 promoters as IRES candidates were co-transfected into HEK293T cells for 24 hours. The diagrams show expected linear transcripts generated from the plasmid CMV promoter (upper, containing red mCherry) and from the Hoxa9 and Chrdl1 promoters (lower, H-GF-Alu and C-GF-Alu). **(D)** Diagram depicts the expected linear mRNAs generated by trans-splicing of Hoxa9 promoter transcripts to CMV promoter transcripts. The resulting chimeric mRNAs have sequences from both the Hoxa9 and Chrdl1 promoter regions. **(E)** RT-PCR experiments detect linear mRNAs containing spliced GFP. Spliced GFP was detected from cDNA generated with RT primers targeting GFP (lanes 1 and 2), and the downstream intron 3’ UTR of trans-spliced transcripts (lanes 3-6). Spliced GFP was also detected in poly-A selected RNA, consistent with the expected linear trans-spliced artifacts. RNase R treatment eliminates the spliced GFP band, confirming the templates were linear RNAs. **(F)** Detection of chimeric products by RT-PCR from total RNA. cDNA generated using the 3’ intron primer were amplified with Hoxa9 (HF) and Chrdl1 (CF) 5’ UTR forward primers and reverse primers specific to the Hoxa9 (HR) and Chrdl1 (CR) 5’ promoter regions. Chimeric (HF / CR and CF / HR) PCR products were amplified from cDNA generated from co-transfected cells (H/C; lanes 3 and 5), but not from control cDNA generated by mixing total RNA from cells transfected with individual Hoxa9 and Chrdl1 promoter plasmids (H+C; lanes 4 and 6). This confirms that chimeric products arise from trans-splicing in HEK293T cells, rather than from template switching during RT-PCR.

“… most circular RNA over-expression constructs still backsplice at low efficiency and generate many undesired transcripts, including unspliced RNAs, concatamers, or *trans-* spliced RNAs (Ho-Xuan et al, 2020; Fig. 5). These undesired RNAs all potentially limit the utility of these constructs for defining circular RNA functions.” (Dodbele *et al*, 2021).

We previously showed that putative *Hoxa9* and *Chrdl1* IRESes contain promoters (Akirtava *et al*, 2022). These promoters could conceivably generate *trans*-spliced linear artifacts in the toolbox study’s reporter plasmid (Fig. 1B). To test this, we recreated the toolbox study’s *Hoxa9* and *Chrdl1* reporter plasmids containing putative IRESes and co-transfected them into HEK293T cells (Fig. 1C). Linear *trans*-spliced artifacts are expected to have spliced GFP followed by the putative IRES, a second copy of the 5’ GFP fragment, and an unspliced intron (Fig. 1D), while circular RNAs should have only the test IRES sequence and GFP. To compare the abundance of linear and circular spliced GFP, we first amplified spliced GFP by RT-PCR using an RT oligo complementary to GFP (Fig. 1E, lane 1). Importantly, RNase R treatment effectively eliminated the spliced GFP band, suggesting spliced GFP was almost completely linear (Fig. 1E, lane 2). We also used a primer specific to the unspliced intron (Alu 1) to reverse transcribe linear reporter RNAs, and PCR amplified spliced GFP (Fig. 1E, lanes 3-6). This readily detected spliced GFP from both total and poly-A selected RNA, the latter of which should be highly depleted of circRNA transcripts, as the predicted circRNA products would lack a poly-A tail. RNase R treatment also eliminated spliced GFP detection, confirming that spliced GFP mRNAs were primarily linear.

To further test the hypothesis that these linear GFP mRNAs were generated by *trans-* splicing, we used PCR oligos targeting pairs of sequences derived from the separate *Hoxa9* and *Chrdl1* transcripts (Fig. 1F). This confirmed that the linear GFP mRNAs were generated by *trans*-splicing (Fig. 1F, lanes 3 and 5). Additional controls established that these chimeric *Hoxa9*/*Chrdl1* RT-PCR products were not generated by RT-PCR strand-switching (Fig. 1F, lanes 4 and 6; McManus *et al*, 2010) and are instead produced by *trans*-splicing. The presence of *trans*-spliced artifacts makes it impossible to determine whether GFP from the back-splicing reporter plasmid is produced by cap-dependent translation of linear *trans*-spliced RNAs or IRES-mediated translation of back-spliced circRNAs. Thus, GFP expression from the proposed IRES “toolbox” plasmid is not a reliable measure of IRES activity.

To quantify the expression of *trans*-spliced linear GFP products, we transfected HEK293T cells with the *Hoxa9*, *Chrdl1*, and *Dlx1* back-splice reporter plasmids for 24 hours and performed RNA-seq with polyA selection. The resulting reads were aligned to both the human genome and transfected plasmids, and GFP splice-junction reads were used to estimate expression levels relative to cellular genes using the RPKM metric (Fig. 2A). Spliced GFP levels were among the most abundant transcripts (1^st^ percentile) in polyA selected RNA-seq for all three reporter constructs (Fig. 2B), with levels similar to highly expressed genes, e.g. RPL18A. We also observed multiple cryptic splicing events generated from the back-splice plasmid, including a cryptic 5’ splice site at the mRuby reporter stop codon (CAA|G**<u>UAA</u>**GA) that spliced to the 3’ GFP fragment with high efficiency (Fig. 2C). To evaluate the extent to which polyA selection removed circRNA from our samples, we sequenced one replicate of each transfection using rRNA depletion. PolyA-selection eliminated back-splice junction reads from the relatively abundant cellular circHIPK3(Zheng *et al*, 2016) and greatly reduced the abundance of non-adenylated histone mRNAs. In contrast, 11-15% of spliced GFP was recovered in the polyA dataset (Fig. 2D; Table S1). These results show back-splicing plasmid *trans*-spliced linear artifacts are expressed at levels sufficient to drive the production of measurable GFP protein in transfected cells.

**Figure 2.**
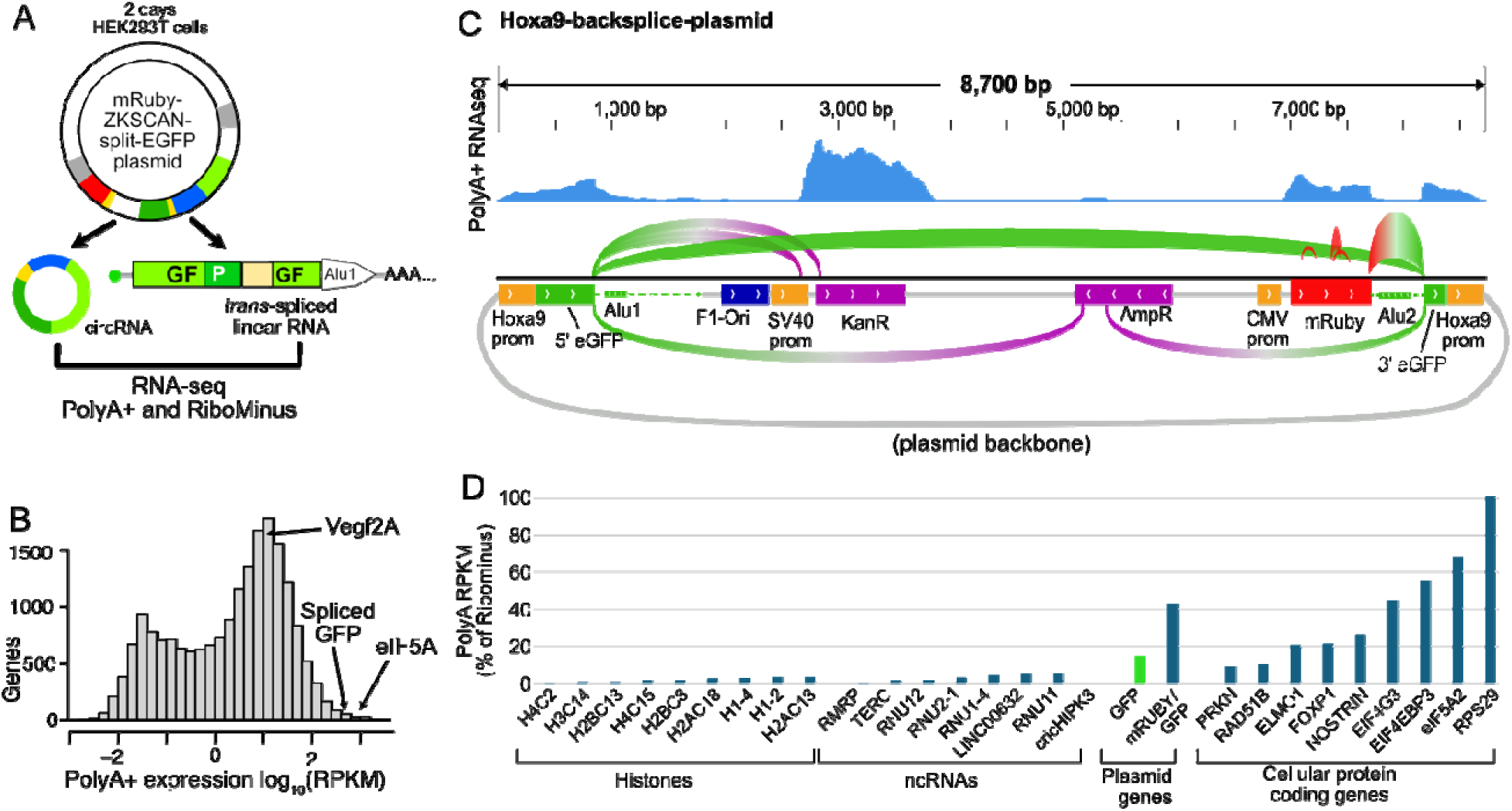
The “toolbox” back-splicing plasmid produces abundant spliced, polyadenylated GFP mRNA and other aberrant products. **A.** The back-splicing plasmid can produce both non-adenylated circRNA (left) and polyadenylated *trans*-spliced linear mRNAs (right). To quantify these, HEK293T cells were transfected with plasmids (two biological replicates each) containing the *Hoxa9*, *Chrdl1*, and *Dlx1* promoters as candidate IRESes and subjected to polyA+ and Ribominus RNA-seq (one sample). **B.** Histogram depicting the average expression level of spliced GFP from the three plasmids compared to cellular transcripts. Spliced, polyadenylated GFP mRNA was expressed in the top 1% of endogenous mRNAs. **C.** Genome browser view depicting polyA+ RNA-seq results from the *Hoxa9* promoter containing back-splicing plasmid. Short-read coverage is shown above, while arcs show the most frequent splice-junctions from the plasmid. The long arc depicts frequent splicing of 5’ and 3’ GFP, consistent with *trans*-spliced mRNAs identified in Figure 1. Additional splicing events occured between 5’ GFP, the reverse complement of the AmpR gene and 3’ GFP, as well as splicing between 5’ GFP and downstream cryptic splice sites in the SV40 promoter and KanR gene. The mRuby mRNA is almost exclusively spliced to the 3’ fragment of GFP. Many other unexpected splicing events were present at lower levels, but not shown for simplicity. **D.** PolyA-selection efficiently depletes non-adenylated mRNAs, including the endogenous circRNA circHIPK3. The average PolyA-selected RPKM values were divided by the RiboMinus RPKM values as a metric of the percentage of expression recovered by poly-A selection. Genes whose transcripts lack polyA tails (Histones and ncRNAs) were drastically de-enriched after polyA-selection, as was the endogenous cellular circRNA circHIPK3 (0%). At least 15% of the spliced GFP expressed from the plasmid contains polyA tails, consistent with *trans*-spliced artifacts. Representative cellular linear protein coding transcripts have a wide range of polyA+ enrichment, suggesting varying levels of deadenylation and / or decay intermediates.

### Relevance of “promoterless” reporter controls in error-prone plasmid IRES assays

Because promoters create monocistronic linear artifacts in both bicistronic (Kozak, 2001; Bert *et al*, 2006; Han & Zhang, 2002; Wang *et al*, 2005; Huang *et al*, 2007) and back-splicing circRNA plasmid IRES assays (Fig. 1; Chu *et al*, 2021; Jiang *et al*, 2021; Ho-Xuan *et al*, 2020; Dodbele *et al*, 2021; Unti *et al*, 2024), tests of promoter activity have been considered key controls in plasmid IRES reporter assays (reviewed by (Thompson, 2012; Terenin *et al*, 2017; Yang & Wang, 2019; Loughran *et al*, 2025)). The toolbox study argued that promoter activities in candidate IRESes are irrelevant under the assumption that their plasmid was unaffected by promoter artifacts. However, our results demonstrate that this plasmid generates linear *trans*-spliced artifacts (Fig. 1). Additionally, the toolbox study implied that promoters in its nominated “IRES” candidates are “cryptic” promoters that are not “active in native chromatin”. This interpretation is contradicted by multiple independent datasets. We and others have shown that the putative IRESes from *Hoxa3, Hoxa5, Hoxa9,* and *Chrdl1* are native promoters, supported by data from *in vivo* experiments including histone modification patterns from native chromatin, nAnT-iCAGE transcription start site (TSS) maps from mouse embryo neural tube, EPDnew promoter annotations, and functional plasmid assays (Ivanov *et al*, 2022; Akirtava *et al*, 2022). Thus, characterizing these promoters as “cryptic” is directly contradicted by multiple lines of evidence.

The toolbox study also misinterpreted the effects of *Hoxa9* promoter mutations when concluding that putative IRES promoter activities are irrelevant. The *Hoxa9* promoter has a binding site for transcription factors USF1 and USF2 supported by mouse and human ChIP-seq data and siRNA depletion experiments (Zhang *et al*, 2020; Akirtava *et al*, 2022). The USF1/2 consensus motif reported in Akirtava et al. is “CA<u>CG</u>TG<u>AC</u>” (Wang *et al*, 2012; Lambert *et al*, 2018). The toolbox study mistook this motif as “CANNTG”, lacking the internal “CG” and 3’ “AC” sequences. This led them to incorrectly conclude their “M13” mutation does not alter the USF1/2 site, even though it changes the site from “CACGTG<u>AC</u>” to “CACGTG<u>TG</u>”. Contrary to the toolbox study’s interpretation, reduction of both RNA and protein expression by M13 (See Figs 2G, 4D, and 4E from (Koch *et al*, 2025)) can be explained by disruption of the USF1/2 binding site in the *Hoxa9* promoter. The toolbox study’s M2 and M5 mutations also disrupt the USF1/2 site and decrease both RNA and protein expression in their reporter plasmids. However, they claimed that M2 and M5 do not decrease RNA levels because the changes are not statistically significant in two-tailed t-tests (p-values = 0.13 for M2 and 0.06 for M5). This false reasoning, that measurements are equal if their differences are not statistically significant, is called the nullification fallacy. It is a logical fallacy because noisy experimental data may lack sufficient power to reject the null hypothesis (that two samples are equal), resulting in a type II error (Harris *et al*, 2012; Kühberger *et al*, 2015). Indeed, the toolbox study’s qRT-PCR data were noisy, likely resulting in false negatives. Furthermore, because the USF1/2 mutations are expected to decrease expression, a one-tailed t-test could arguably be adequate. In that case, the reduced RNA expression could be considered statistically significant for M5 (p = 0.032) and nearly significant for M2 (p = 0.065). Rather than showing promoter controls are irrelevant, the fact that all of these “IRES” mutations in the USF1/2 site of the *Hoxa9* promoter decreased RNA levels underscores the importance of testing for promoter false positives in plasmid IRES assays that are known to be susceptible to such artifacts.

### Orthogonal assays find no IRES activity from *Hoxa9*, *Chrdl1*, and *Dlx1* promoters

Since the back-splicing plasmid used by the toolbox study creates abundant *trans*-spliced linear artifacts (Figs. 1 & 2), we sought to validate three putative IRES elements they reported from *Hoxa9*, *Chrdl1*, and *Dlx1* promoters using two orthogonal assays (Figs. 3 & 4). These promoter sequences were reported to have IRES activity exceeding that of the EMCV and, in some cases, the HCV viral IRESes (Fig. 3B) in HEK293T cells. We cloned these promoters and the putative *CDK11a* cellular IRES (Cornelis *et al*, 2000) into the Tornado circRNA plasmid between the LgBiT and SmBiT fragments of nanoluciferase. This plasmid generates circRNAs encoding nanoluciferase by cleavage of two Twister ribozymes and circularization by the tRNA ligase RtcB (Fig. 3C). Importantly, the Tornado plasmid was shown to have negligible linear artifacts, and known false-positive IRESes had little activity in this plasmid (Unti *et al*, 2024). We used three viral IRESes (HCV, EMCV, and CrPV) as positive controls, and three randomly scrambled sequences and the 82-nt *Hoxa9* 5’ UTR as negative IRES controls. In the Tornado system, the HCV IRES produced at least 1,400-fold more luciferase than each of the putative cellular IRESes (Fig. 3D; Table S3). Even the CrPV IRES, which originates from an insect virus and has extremely weak activity in human cells, was at least 5-fold more active than any cellular IRES candidate. Sequencing cDNA from transfected cells confirmed that correctly ligated circRNAs were generated from plasmids encoding the putative cellular IRES candidates (Fig. EV2). While the *Chrdl1* and *Hoxa9* promoters made slightly more luciferase than negative controls, this may reflect cap-dependent translation of rare linear artifacts from trans-ligation events (see discussion). These results show three candidate IRESes from the toolbox study have negligible IRES activity in an orthogonal circRNA plasmid that minimizes linear artifacts.

**Figure 3.**
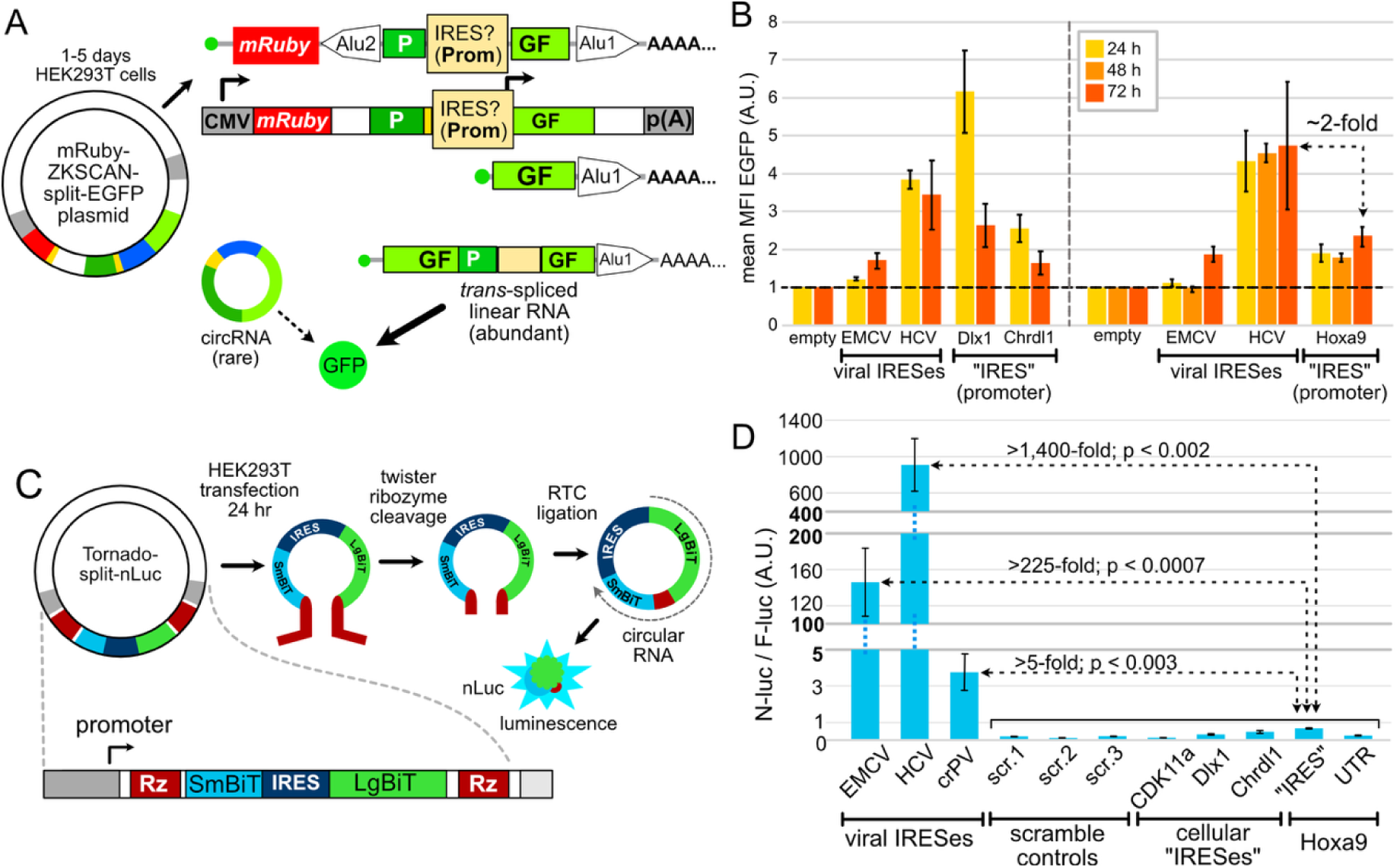
Putative IRESes from Hoxa9 and cellular mRNAs have negligible activity in a ribozyme-based circRNA reporter plasmid. **(A)** Diagram of the ZKSCAN1 plasmid designed to generate circRNA by backsplicing in the toolbox study Promoter activities in putative IRESes produce capped linear transcripts via trans-splicing (see Fig. 1). **(B)** Data from the toolbox study In the ZKSCAN1 back-splicing plasmid, putative cellular IRES elements produced more GFP than the EMCV IRES and half as much GFP as the HCV IRES **(C)** Diagram of the Tornado-split nLuc reporter system. The Tornado plasmid generates circRNA via ribozyme cleavage and ligation by RtcB, resulting in extremely low levels of linear RNAs. **(D)** Putative IRES elements from CDK11a, Dlx1, Chrdl1, and Hoxa9, the 82-nt Hoxa9 5’ UTR from NCBI Refseq annotations, and negative control sequences (randomly scrambled CDK11a) were inserted between LgBit and SmBiT. The putative IRES elements have negligible activity in the Tornado circRNA system (n = 6; compare to B). The Y-axis has two breaks, which were necessary to show the rare signal from putative cellular IRESes in this experiment.

We also tested these putative cellular IRESes using direct RNA transfection, which avoids problems caused by plasmid-based promoter artifacts (Terenin *et al*, 2017; Loughran *et al*, 2025). In our RNA transfection assay, candidate IRES mRNAs were constructed with either an m7-G cap (positive control) or an A cap with a stable 5’ stemloop to inhibit directional scanning (Russell *et al*, 2023; Kozak, 1989; Fig. 4A). As with the Tornado plasmid, direct RNA transfection with the A cap / stemloop RNAs showed negligible activity from any of the candidate IRES elements as compared to HCV, equivalent to random sequence negative controls (Fig. 4B). Notably, the genuine 82-nt *Hoxa9* 5’ UTR (Akirtava *et al*, 2022; Ivanov *et al*, 2022), which lacks previously reported critical “IRES” sequences, drove as much translation from A-cap / stemloop mRNAs as the putative cellular IRESes. m7-G capped mRNAs had much higher translation, as expected from cap-dependent translation initiation. Indeed, the m7-G capped bona fide *Hoxa9* 5’ UTR produced 1,000-fold more luciferase than the putative *Hoxa9* A-cap / stemloop IRES construct (Fig. 4B; Table S3; Fig. EV3). These results show that putative IRES elements nominated by the toolbox study using the error-prone back-splicing plasmid cannot be validated by orthogonal methods, and that cap-dependent translation is at least three orders of magnitude more efficient than cap-independent translation from these sequences.

**Figure 4.**
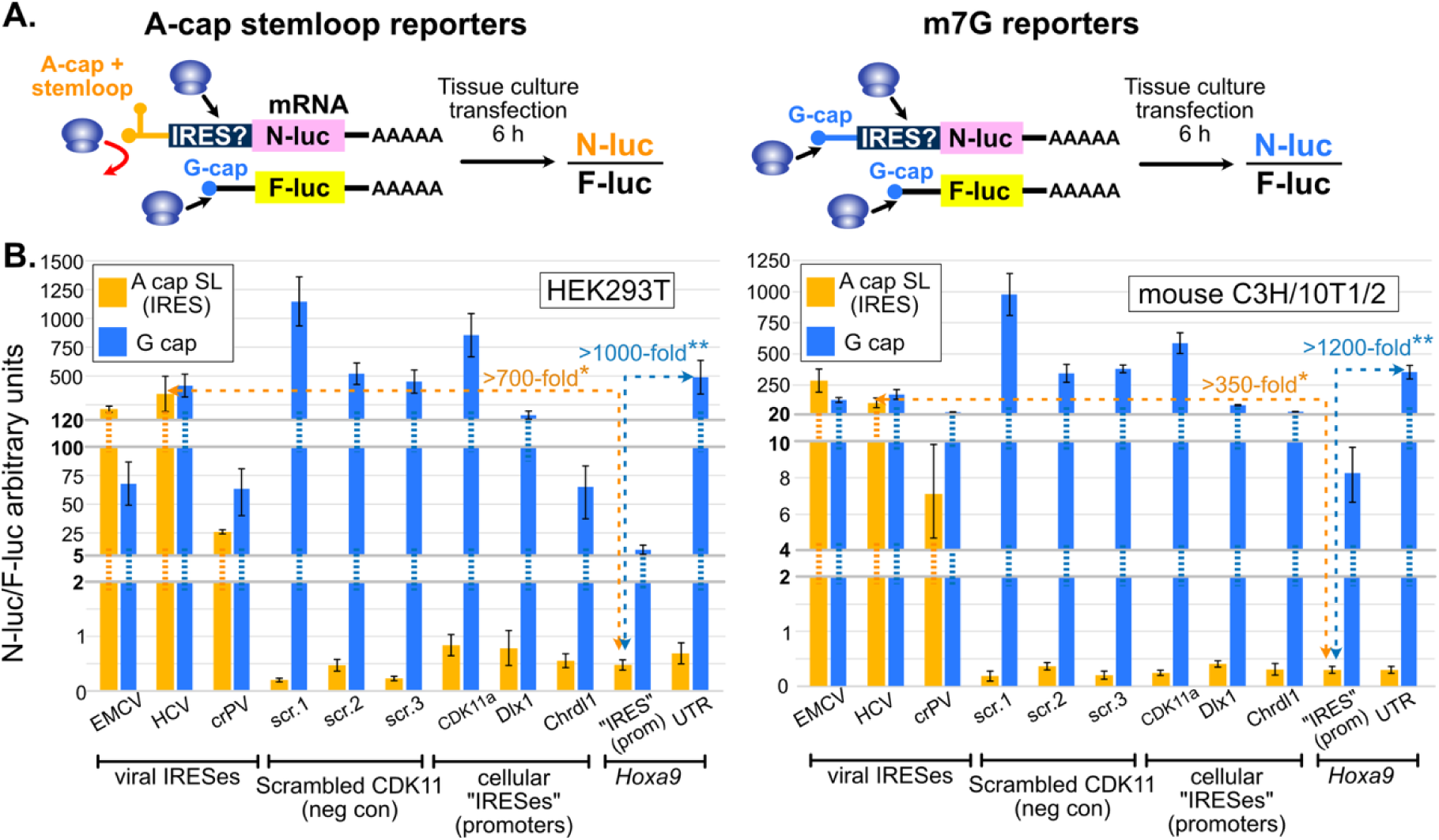
Putative cellular IRESes have negligible “IRES” activity in mRNA transfection assays. **(A)** 5’ UTRs were cloned upstream of luciferase reporters to compare expression from IRES (orange) and m7G-cap scanning (blue) mechanisms. IRES reporter mRNA constructs greatly reduce cap-dependent scanning by using an adenosine “A” cap followed by a strong stemloop. Viral IRESes from EMCV, HCV, and CrPV were used as positive controls. Negative controls were randomly scrambled CDK11a (see Fig. 3). IRES and G-cap mRNAs were co-transfected with m7G-capped firefly luciferase for normalization **(B)** Viral IRESes expressed similar levels of Nluc in IRES and G-cap reporters. In contrast, the putative cellular IRESes all exhibit IRES activity equivalent to scrambled negative control sequences. The genuine 85 nucleotide Hoxa9 5’ UTR, which lacks all previously reported “IRES” essential sequences, has slightly more IRES activity than the Hoxa9 promoter region claimed to be an “IRES” by the toolbox study G-cap transcripts with the genuine Hoxa9 5’ UTR produced >1000-fold more luciferase than the Hoxa9 “IRES”. Similarly, G-cap Chrdl1 and Dlx1 reporters produce ∼100-fold and ∼200-fold more luciferase than corresponding “IRES” reporters. The Y-axis has two breaks, which were necessary to show the weak signal from putative cellular IRESes and negative controls. (n=4; * = p < 0.05; ** = p < 0.01; Also see Table S3 and Fig. EV3).

### PacBio Iso-Seq data are supported by transcription start sites from an orthogonal approach

Identification of bona fide transcript 5’ ends is necessary for IRES studies to ensure candidate IRESes are part of legitimate 5’ UTRs. PacBio Iso-Seq uses oligo-dT primers for reverse transcription and template switching oligos to map the 3’ and 5’ ends of full-length cDNA, respectively (Fig. 5A). The toolbox study used PacBio Iso-Seq to sequence mRNA transcripts from mouse embryonic somites and neural tube tissue. Upon observing PacBio 5’ UTRs that were shorter than several transcripts annotated in GENCODE, the toolbox study concluded that 5’ UTRs are frequently truncated in PacBio Iso-Seq reads and cannot be trusted. Contrary to this conclusion, other studies have reported that 80% to 96.5% of PacBio Iso-Seq transcription start sites (TSSs) correspond to TSSs mapped using CAGE (Wen *et al*, 2023; Wyman *et al*, 2019; Leung *et al*, 2021). Furthermore, PacBio sequencing of *in vitro* transcribed control RNAs showed 5’ truncation only begins to be an issue for RNAs 6 kb or longer (ext. data Fig. 3 in Pardo-Palacios *et al*, 2024b). Since nearly all mouse mRNAs are less than 4 kb long (Bussotti *et al*, 2016; Adams & Vollmers, 2024), 5’ truncation is most likely not a common problem in PacBio Iso-Seq data. Instead, the toolbox study’s chosen GENCODE annotations may include erroneous transcript models with incorrect extended 5’ UTRs and may be missing genuine transcripts with shorter 5’ UTRs. This could be evaluated by comparing their PacBio Iso-Seq 5’ ends to TSSs mapped using an orthogonal method in the same biological context.

**Figure 5.**
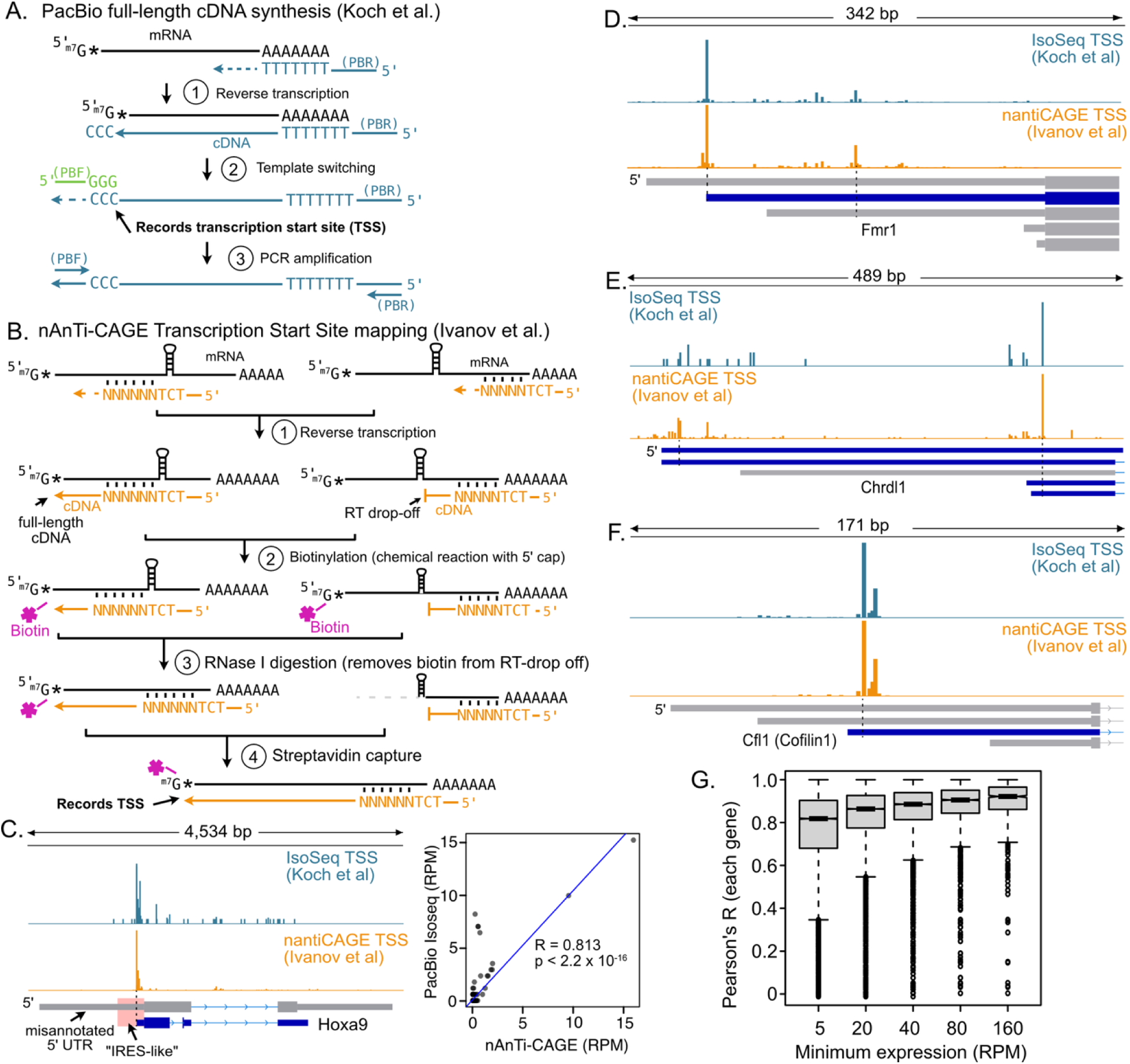
PacBio Iso-Seq maps accurate transcription start sites (TSSs). **(A)** Full-length cDNA cloning in PacBio HiFi Iso-Seq. An oligo-dT primer (blue) is used for first-strand cDNA synthesis. Reverse transcriptase (RT) adds three untemplated “C” nucleotides at the cDNA 3’ end at the m7G cap. This records the TSS location. RT then switches templates to use an oligo with “GGG”-3’ (green) to complete first-strand synthesis. PacBio specific forward (PBF) and reverse (PBR) primers are used to PCR amplify the cDNA library. **(B)** Diagram of nAnTi-CAGE. Random hexamers (blue) are used for first strand cDNA synthesis, allowing bypass of structures that could cause RT drop-off. The m7G cap is biotinylated through a chemical reaction. RNase I removes the biotinylated 5’ ends of any RNAs with premature RT drop-off. The mRNA / cDNA hybrids are recovered by streptavidin affinity capture, and the cDNA 5’ ends are sequenced. **(C)** IGV Genome browser snapshot of Iso-Seq (toolbox study; blue) and nAnTi-CAGE read 5’ ends (Ivanov et al.; orange) from E11.5 mouse embryonic somites and neural tubes. The putative Hoxa9 “IRES isoform” annotated by GENCODE is shown in gray, and the “IRES-like” element is highlighted in pink. There is no evidence of a TSS corresponding to the “IRES isoform”. The inset graph compares the normalized number (RPM) of 5’ ends mapped by PacBio Iso-Seq and nAnTi-CAGE in a 1kb region around the transcription start site depicted in the blue transcript model. **(D-F)** Genome browser snapshots comparing the distribution of 5’ end reads from these orthogonal datasets for *Fmr1*, *Chrdl1*, and *Cofilin1*. TSSs for these genes are nearly identical in the two datasets, indicating PacBio Iso-Seq is not subject to extensive RT drop-off. GENCODE 5’ UTRs supported by PacBio and nAnTi-CAGE TSSs in these tissues are shown in blue, while unsupported isoforms are shown in gray. **(G)** Boxplot of the distributions of Pearson’s R values for the TSS frequencies at single nucleotides in 1kb windows around transcript isoforms genome wide. The two datasets are highly concordant.

In this case, a genome-wide TSS dataset from an orthogonal method is available. Ivanov et al., 2022 mapped TSSs in the same tissue and embryonic stage using no-amplification non-tagging CAGE (nAnT-iCAGE) (Ivanov *et al*, 2022). This method uses random priming for cDNA synthesis, chemical biotinylation of 5’ m7G caps, RNase I digestion to remove biotin from potentially truncated cDNA “drop off” products, streptavidin capture, and ligation-based Illumina sequencing (Fig. 5B) (Murata *et al*, 2014). By using random primers, nAnT-iCAGE initiates reverse transcription at multiple locations, which bypasses RNA structures that might otherwise cause premature cDNA drop-off (Fig. 5B). Combined with RNase I digestion, this approach ensures that only completely synthesized cDNA from 5’ capped RNAs are sequenced (Cvetesic *et al*, 2018). The nAnTi-CAGE TSS data from Ivanov et al. are thus ideally suited to evaluate the frequency of RT drop-off in the PacBio Iso-Seq reads from the toolbox study.

We first compared the number of 5’ end reads in the toolbox study and Ivanov et al. datasets at each nucleotide around genes which the toolbox study interpreted as having truncated 5’ UTRs. For example, the frequency of 5’ ends mapped at each nucleotide by the two methods were highly correlated for the *Hoxa9* gene (Fig. 5C), such that neither method supports the misannotated 5’ UTR containing the putative *Hoxa9* IRES. We found similar high correlations genome wide, examples of which are shown for *Fmr1*, *Chrdl1*, and *Cfl1* (Fig. 5D-F). Strong positive correlations were observed for the vast majority of transcripts and were stronger with increasing expression (median Pearson’s R ranging from 0.8 to 0.9; Fig. 5G). These results show that both the location and relative frequencies of PacBio Iso-Seq 5’ ends are remarkably consistent with 5’ ends identified by nAnT-iCAGE, an orthogonal technology that is not susceptible to RT drop-off. Thus, the claim that these PacBio 5’ ends are inaccurate is contradicted by an orthogonal dataset from matched tissue. We conclude that the toolbox study’s PacBio 5’ ends are overwhelmingly supported as TSSs and can be trusted for accurate mRNA transcript annotation.

### Reanalysis of PacBio Iso-Seq data shows the *Hoxa9* promoter region (“IRES” isoform) is transcribed only as non-coding RNA

We and others previously reported that the long 1,266 nucleotide (nt) 5’ UTR isoform of mouse *Hoxa9* (“IRES isoform”), containing the *Hoxa9* promoter, was misannotated, and that the 5’ UTR is only 82 nt long (Akirtava *et al*, 2022; Ivanov *et al*, 2022; Fig. EV1). The data supporting this include short-read (Illumina) RNA-seq from mouse embryos and tissues, annotated transcription start sites from refTSS (Abugessaisa *et al*, 2019), annotated promoter elements from EPDnew (Dreos *et al*, 2015), ENCODE promoter elements based on DNase I hypersensitivity and H3K4me3 (ENCODE Project Consortium *et al*, 2020), paused RNAPII ChIP-seq peaks (Eng *et al*, 2014), nAnT-iCAGE TSS maps from mouse embryonic neural tube and somites (Ivanov *et al*, 2022), and public PacBio Iso-Seq data from mouse embryonic forelimbs. The extended “IRES” isoform is also much longer than the *Hoxa9* mRNA length reported by northern blotting in prior studies (Akirtava *et al*, 2022; Ivanov *et al*, 2022). PacBio Iso-Seq reads from mouse embryo forelimbs also indicated the presence of non-coding *Hoxa10*/*Hoxa9* and *Mir196b*/*Hoxa9* fusion transcripts generated by run-on *Hoxa10* transcription through the *Hoxa9* promoter region and splicing to *Hoxa9* exons (Akirtava *et al*, 2022; Fig. EV1). The combination of overlapping coding and non-coding transcripts expressed from the *Hoxa9* locus makes it a challenging genomic region to annotate.

To further evaluate mRNA isoforms expressed during development, the toolbox study generated PacBio Iso-Seq data from E11.5 somites and neural tube. An alignment of these PacBio reads is shown in Fig. 6, which corresponds to toolbox study Fig. EV1. In both our and their alignment figures, no reads match the complete proposed “IRES” isoform of *Hoxa9* (Fig. 6; compare to Fig. EV1 from the toolbox study). In addition, several reads correspond to *Hoxa10*/*Hoxa9* and *Mir196b*/*Hoxa9* fusion transcripts, precursors for mir-196b (pri-mir196b transcripts) (Fig. 6, compare to Fig. EV1 and Fig. 6A from the toolbox study). Instead of reporting what is visible in their alignment, the toolbox study assigned transcript isoforms to their data using a program called Isoquant (Prjibelski *et al*, 2023). Isoquant is sensitive to user input parameters because it first assigns PacBio reads to transcript annotations provided by the user. Reads that cannot be assigned to the user-provided annotations are then used to predict novel transcripts. In this way, Isoquant relies heavily on the gene annotation provided by the user (Pardo-Palacios *et al*, 2024b). The toolbox study provided Isoquant with two predicted *Hoxa9* isoforms, “A” (3,226 nt long) and “B” (851 nt long), from the GENCODE M32 annotation. Notably, these isoforms are inconsistent with published datasets (Akirtava *et al*, 2022) and with the previously reported length of *Hoxa9* mRNA based on northern blotting from embryos (1,900 - 2,500 nt)(Rubin *et al*, 1987; Fujimoto *et al*, 1998), especially considering an average polyA tail length of 250 nucleotides (Kühn *et al*, 2009). Isoform A has an incorrect, extended 5’ UTR (Fig. 6A). Isoform B has an incorrect truncated 3’ UTR (Fig. 6A). Provided with these two incorrect options, Isoquant assigned PacBio reads primarily to Isoform A. Thus, the toolbox study provided Isoquant with incorrect annotations and then reported that the program detected the misannotated “IRES” isoform. Visual analysis of the read alignments from the toolbox study argues against this isoform, suggesting incorrect annotations allowed Isoquant to choose one incorrect isoform over another incorrect isoform as a better match to the raw data.

**Figure 6.**
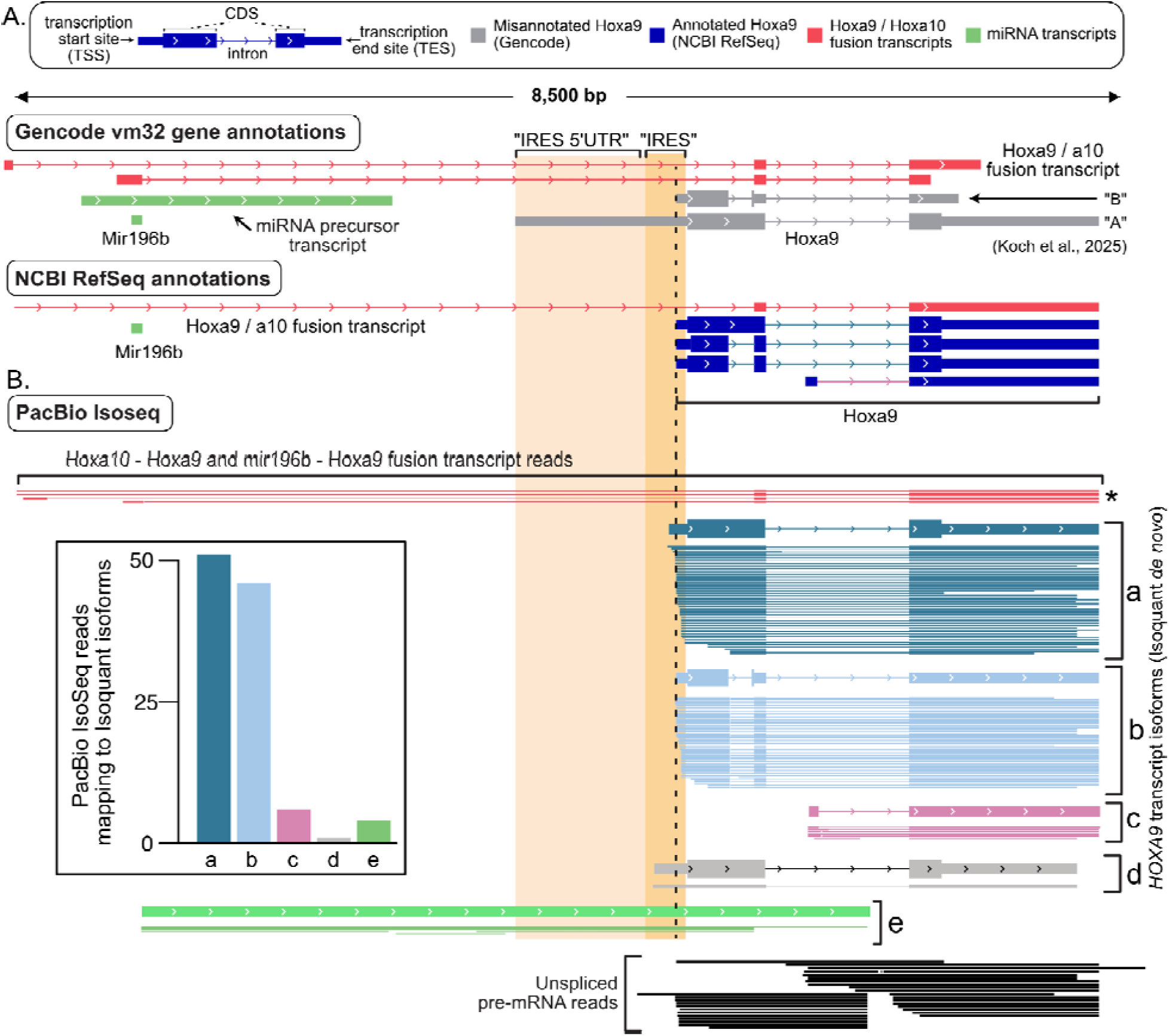
Putative Hoxa9 IRES isoform corresponds to non-coding RNA in mouse somites and neural tube. **(A)** Genome browser snapshot with transcript models from GENCODE and NCBI. The GENCODE M32 annotation used by the toolbox study includes two misannotated Hoxa9 isoforms – “A” with an extended 5’ UTR (gray) and “B” with a truncated 3’ UTR. The putative “IRES-like” and “5’ UTR” regions of the “A” isoform are shaded for comparison. Three precursor transcripts for mir196b are also annotated, including two fusion transcripts between mir196b and Hoxa9. **(B)** Alignment of PacBio Iso-Seq data from the toolbox study are shown. Four PacBio reads from spliced Hoxa10 / Hoxa9 and mir196b / Hoxa9 fusion transcripts are depicted in red (asterisk). After removing Hoxa9 locus transcripts from GENCODE M32, Isoquant predicted five transcripts in the Hoxa9 region (labelled a-e). The supporting PacBio HiFi reads are shown below each transcript. The inset bar graph indicates the number of reads corresponding to each Isoquant transcript. Transcripts a and b (mauve and dark blue) are supported by 93% of the PacBio HiFi reads. No HiFi reads correspond to the fully extended putative IRES 5’ UTR isoform (“A”). Transcript model e (green) is consistent with the F3 product of miR196b processing from primary transcripts. Reads from unspliced Hoxa9 pre-mRNA are shown below (black). One (0.96%) read covers most of the putative IRES region in transcript d. The misannotated IRES transcript (gray) completely overlaps the non-coding pri-miR196b and non-coding fusion transcripts introns.

To directly assess how the provided annotations might have affected their Isoquant results, we removed the *Hoxa10*, *Mir196b*, and *Hoxa9* gene annotations and reran Isoquant with the toolbox study’s PacBio data. This approach allowed Isoquant to assemble *Hoxa9* locus transcripts *de novo*, such that it defined five transcript isoforms which we designate a, b, c, d, and e (Fig. 6). The 5’ ends of all these isoforms are supported by nAnT-iCAGE data (Fig. 5) and the isoforms closely match NCBI RefSeq annotations (Fig. 6). Importantly none of these isoforms correspond to the 3,226+ nt putative *Hoxa9* “IRES” isoform (Isoform “A” in the toolbox study). Our isoforms a (2,105 nt) and b (1,872 nt) represent 93% of PacBio *Hoxa9* reads and differ by the splicing of an alternative intron. These isoforms both have the short 5’ UTR and the extended 3’ UTR, consistent with northern blot estimates of *Hoxa9* mRNA length (1,900-2,500 nt; (Rubin *et al*, 1987; Fujimoto *et al*, 1998)). Isoform c is rare (∼6%), and initiates transcription within the large intron shared by isoforms a and b. Isoform d has one read (0.96%) extending 121 nt upstream of the major TSS, into the putative “IRES” region. Isoform e is particularly interesting, as its 5’ end corresponds precisely to the 3’ end of mir196b and it extends well into the *Hoxa9* mRNA coding sequence (CDS). This isoform is consistent with an expected downstream product of Drosha cleavage of the primary miR196b transcript (Fig. EV4; Kim *et al*, 2025). In addition, the reads supporting isoform e extend completely through the putative “IRES” isoform region indicating reverse transcriptase can fully traverse this RNA without premature “drop off”. Notably, multiple reads from the toolbox study show fusion transcripts between *Hoxa9* and upstream genes. This finding from their provided data contradicts their claim that *Hoxa10*/*Hoxa9* fusion transcripts are not present. In summary, the toolbox study’s PacBio data show no evidence of a 3,226+ nt *Hoxa9* “IRES” messenger RNA. The toolbox study claimed that this is due to premature 5’ truncation of PacBio reads, which is incorrect (Fig. 5). Instead, the toolbox study data validate the documented short 5’ UTR of *Hoxa9* (Isoforms a and b) (Ivanov *et al*, 2022; Akirtava *et al*, 2022; Adams & Vollmers, 2024) and support the presence of previously reported fusion transcripts, a miRNA processing product containing the “IRES” region, and intronic RNAs excised from fusion transcripts, the latter two of which include the *Hoxa9* promoter sequence and are predicted to be retained in the nucleus.

### ncRNA from the *Hoxa9* promoter (“IRES”) localizes to the nucleus in mouse embryos

To evaluate the expression of *Hoxa9* locus transcripts in a physiological context, the toolbox study performed smFISH using one probe to the putative IRES region and one to the *Hoxa9* coding region (CDS; See probe design in Fig. 7A). However, our analysis indicated that the primary *Hoxa9* mRNA does not include the “IRES” probe target sequence (Fig. 6), but that the “IRES” probe would instead detect the non-coding pri-mir196b transcript and excised introns (Fig. 7A). Additionally, the CDS probe could detect three RNA species – non-coding pri-mir196b transcripts, excised introns, and the *Hoxa9* mRNA (Fig. 7A). Thus, the smFISH probes used by the toolbox study are not specific for potential “IRES” containing mRNA isoforms, as they target non-coding RNAs like pri-mir196b and fusion transcript introns that are expected to be retained in the nucleus. Strikingly, we reanalyzed the images from the toolbox study and found 96% of the “IRES” probe signal in their published images colocalizes with DAPI-stained nuclei in somites and, more rarely, in neural tube regions (Fig. 7B). If the “IRES” was part of a mRNA it would have been abundant in the cytoplasm. Yet it is overwhelmingly nuclear in the images published by the toolbox study. Signal from the “CDS” probe is also primarily nuclear in somites and both nuclear and cytoplasmic in neural tube (Fig. 7B). This is not surprising, as this probe also targets pri-mir196b and unspliced *Hoxa9* pre-mRNA transcripts. Notably, prior work found the mir196 family of miRNAs is specifically expressed in the same region of mouse E10.5-12.5 embryos, posterior somites and neural tube (Wong *et al*, 2015), in which the toolbox study reported signal from their smFISH “IRES” probe. Thus, the smFISH data presented by the toolbox study are inconsistent with the *Hoxa9* “IRES” region being expressed in a cytoplasmic mRNA available for translation by the ribosome. Instead, their results are consistent with the expression of non-coding transcripts that can be observed in their PacBio Iso-Seq data (Fig. 6).

**Figure 7.**
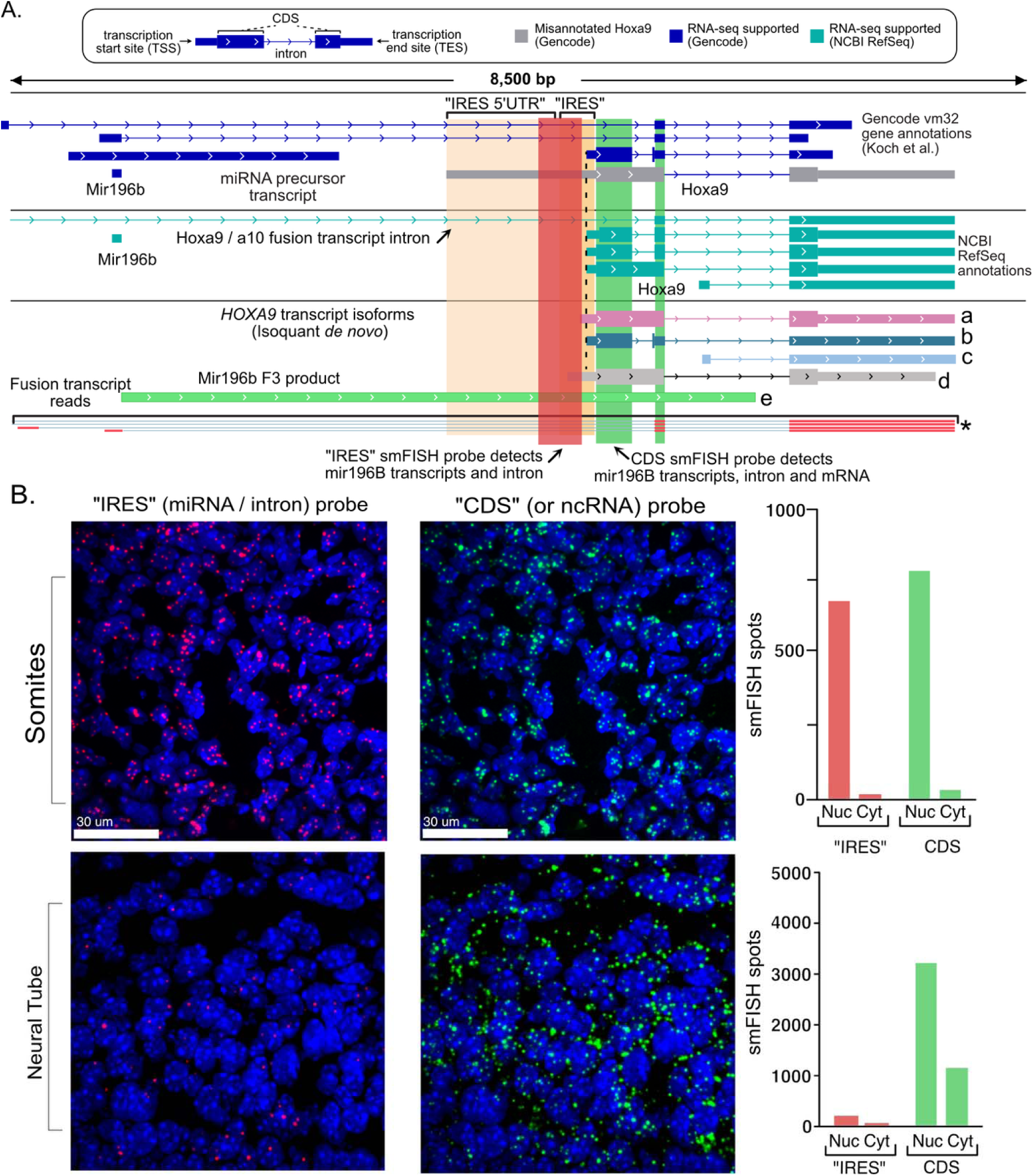
The Hoxa9 “IRES” probe targets ncRNA and is restricted to the nucleus in mouse somites and neural tube. **(A)** Genome Browser diagram shows the smFISH probe locations used in the toolbox studyKoch et al. GENCODE and RefSeq gene annotations are shown, including the misannotated Hoxa9 isoform (gray). The five Isoquant predicted transcripts (a-e) are shown. Four fusion transcript reads are shown (red, asterisk). The “IRES” probe (red) detects the non-coding pri-mir196b transcript (e) and introns from fusion transcripts. The CDS probe also detects these non-coding RNAs, but can additionally detect the mature Hoxa9 mRNA. **(B)** DAPI (blue) and smFISH images from the toolbox study. Fig. 3D were overlayed to visually compare the nuclear localization of “IRES” (red) and “CDS” (green) probe spots. The DAPI nuclear signal shows nuclei with bright heterochromatic regions called chromocenters (Probst & Almouzni, 2011). Bar graphs show quantification of red and green probe spots in nucleus and cytoplasm from the published images.

**Figure 8.**
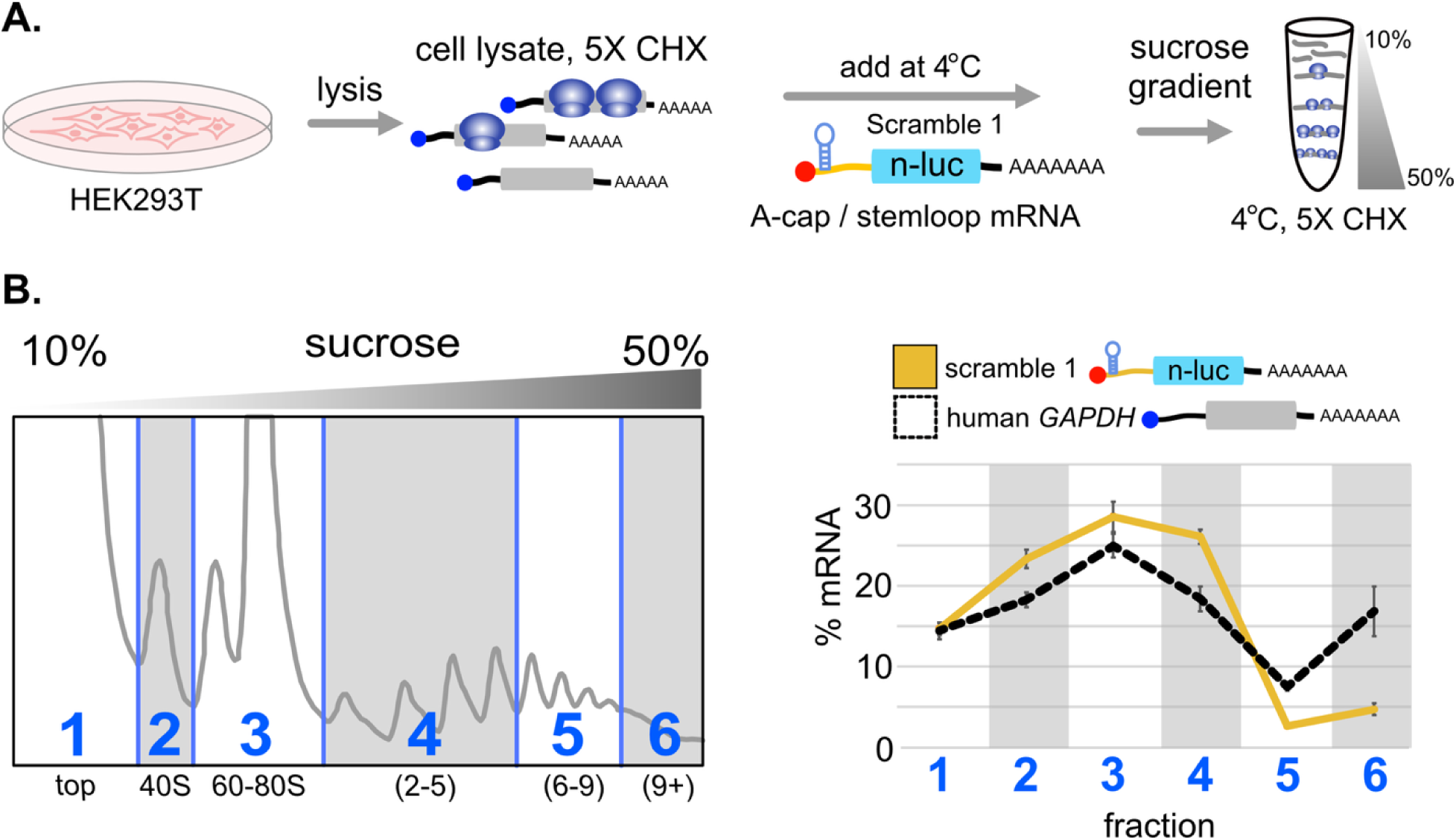
Contaminating, non-translating RNAs can migrate in polysomal fractions. **(A)** A-cap / stemloop IRES reporter RNA carrying the negative control “scramble 1” sequence was mixed into a polysome lysate made from HEK293T cells for five minutes on ice and loaded on a sucrose gradient. **(B)** The sucrose gradient was fractionated and cDNA was prepared from groups of fractions as shown (numbered 1-6). qRT-PCR (n=4) was performed to quantify the exogenous “scramble 1” RNA across the fractions, as compared to endogenous Gapdh mRNA. The “scramble 1” transcript migrated into polysomal fractions 4-6, even though it is not translated (Fig. 4).

### Association of non-translating RNAs in polysome fractions

The toolbox study also used qRT-PCR to detect *Hoxa9* locus transcripts associated with translating ribosomes in deep polysome fractions from mouse embryos. Unlike PacBio Iso-Seq, which is specific for 5’ capped, polyadenylated messenger RNA, qRT-PCR can amplify any RNA containing sequences matching the primers. The “IRES” and “CDS” PCR primers used by the toolbox study are complementary to pri-mir196b, fusion transcripts, and fusion transcript introns. The targeted amplicons are present in every one of these RNA species. Given that smFISH data show RNAs that include the “IRES” region are retained in the nucleus (Fig. 7), PCR amplification of these RNAs from polysome extracts could reflect contamination of polysome extracts with nuclear pri-mir196b ncRNA. Indeed, previous work has found non-coding RNAs associated with polysomes (Fonouni-Farde *et al*, 2021; Bazin *et al*, 2017; van Heesch *et al*, 2014; Yoon *et al*, 2012). Some of these non-translating RNAs may have regulatory functions, while others could be contamination resulting from the scaffolding properties of RNA (Fernandes & Buchan, 2021). Indeed, recent work identified mRNAs in translation-initiation-inhibited condensates (TIICs) in yeast cells that migrate in dense polysome fractions even though they are not translating (Glauninger *et al*, 2025)).

Since direct transfection of scramble control A-cap stemloop reporter mRNAs failed to show significant translation, we reasoned these mRNAs could be used to further test the hypothesis that non-translating mRNAs can migrate in polysomal fractions. We prepared a cytoplasmic polysomal extract from untransfected HEK293T cells. We then mixed *in vitro* transcribed non-translating “Scramble 1” IRES negative control mRNA into the polysomal extract on ice and separated polysomes by sucrose gradient fractionation (see methods). Using qRT-PCR, we quantified the distribution of “Scramble 1” across the gradient and compared it to endogenous *GAPDH* mRNA (Fig. 7). The non-translating RNA migrated in polysomal fractions, even though this RNA was not transfected into tissue culture cells in this experiment. These results show that non-translating RNAs can migrate in dense sucrose gradient fractions, most likely due to molecular interactions with other polysomal components that may be stabilized by the temperature and composition of sucrose gradients (4°C, 140 mM KCl, 15 mM MgCl_2_). It may be possible to distinguish translating and non-translating mRNAs that migrate in dense polysome fractions by varying buffer concentrations or using puromycin to release translating ribosomes. However, this may not be effective at removing non-translating RNAs that are polysome-associated by basepairing with translating mRNA or rRNAs.

## Discussion

Since the discovery of viral IRES elements, researchers have sought to find IRES activity in host 5’ UTRs, termed “cellular IRESes”. While cellular IRESes may exist, the plasmid assays most frequently used to nominate candidate IRESes are not reliable. Both bicistronic and back-splicing plasmid assays are subject to false-positive IRES signals due to promoters in candidate IRESes that produce linear transcripts available for cap-dependent translation initiation. The problem of promoter-driven false-positives is made worse by the fact that many 5’ UTR annotations erroneously include genes’ natural promoters, and many genes have multiple promoters within alternative 5’ UTRs (Akirtava *et al*, 2022). Because of this, it is particularly important both to have accurate 5’ UTR annotations and IRES assays that are not subject to false positives. The toolbox study described methods to annotate mRNA and assay IRES activity which were propose as a toolbox for IRES identification. By reanalyzing the toolbox study data and performing additional experiments, we find the conclusions of the toolbox study are unfounded. Each individual tool in the proposed “toolbox” has confounding variables or artifacts and thus carries the risk of nominating more false-positive IRESes. Our results suggest an alternative, robust platform for transcript annotation and IRES identification (Fig. 9).

**Figure 9.**
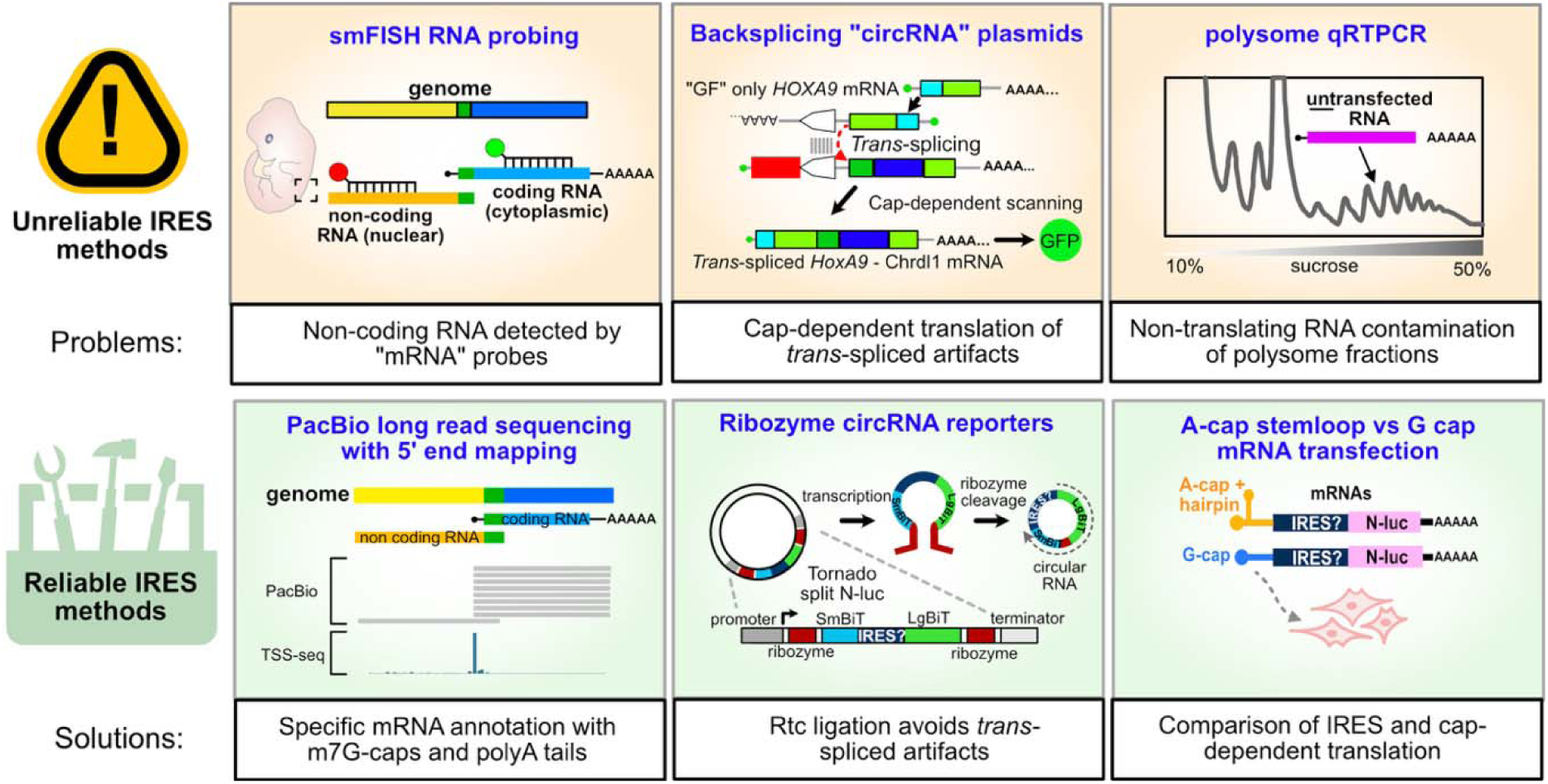
Problems with previously proposed IRES methods and reliable alternatives. Our results show smFISH data can be misinterpreted when using probes that target non-coding RNA, especially when nuclear localization is not considered. Instead, we find PacBio Iso-Seq data provides reliable annotations when analyzed carefully and validated with orthogonal transcription start site mapping data. We also show that the ZKSCAN1 backsplicing plasmid (Chen et al, 2021) used by the toolbox study creates linear artifacts via promoter-driven trans-splicing that manifest as false-positives IRES signals. Instead, we show that the Tornado plasmid system (Unti et al, 2024) is highly resistant to promoter false positives and provides accurate measures of IRES activity when used with appropriate positive and negative controls. Finally, we show that non-translating RNAs can sediment in polysome fractions, which can create artifacts when interpreting polysome association experiments. We instead advocate for direct transfection of IRES reporter mRNAs, which additionally allow direct comparison of the efficiency of IRES-mediated and cap-dependent translation initiation.

First, we provide a simple co-transfection strategy to detect the presence of *trans*-spliced artifacts from back-splicing circRNA plasmids, similar to a previous study (Chu *et al*, 2021). This assay revealed *trans*-spliced linear artifacts produced by the toolbox study plasmids. Indeed, nearly all the spliced GFP generated after 24 hours of transfection was linear, as RNase R eliminated RT-PCR products. Although other work reported that northern blotting could not detect linear artifacts from this plasmid (Figs. S1 and from (Chen *et al*, 2021)(Koch *et al*, 2026)), these experiments used substantially different conditions from the functional assays in the toolbox study. For northern blotting, the back-splice plasmid was diluted 500-fold such that only one in ten cells were expected to receive a single plasmid copy, and RNA was harvested five days after transfection. By that point, RNA decay may deplete linear artifacts, relative to circRNAs, while GFP translated from linear artifacts will have accumulated. Controlling for linear artifacts requires testing each “active” candidate IRES at several timepoints (e.g. 24, 48, and 72 hours). Furthermore, *trans*-spliced linear artifacts would need to be at least 10% as abundant as the circRNA to be detected using the fluorescent probes in such northern blots (Miller *et al*, 2018). Trace linear artifacts (e.g. ∼1% of circRNA) could produce false-positive “IRES” signals not detectable by fluorescent northern blot. Notably, a previous study using radioactive probes for northern blots found a ladder of multiple abundant linear artifacts from a highly related ZKSCAN1 circRNA construct (Ho-Xuan *et al*, 2020). The toolbox study used RNase R treatment and qPCR to demonstrate some exonuclease resistant circRNA was produced after 5 days of transfection, but this strategy assumed that the back-splicing plasmid did not generate *trans-*spliced linear GFP artifacts. It did not examine the presence or absence of linear GFP mRNAs or their levels (Fig. EV5). By targeting the specific structure of *trans*-spliced artifacts, our co-transfection RT-PCR strategy provides a simple, robust alternative to detect false-positives from back-splicing plasmids. However, we see no reason to use the artifact-prone back-splicing reporters for IRES studies, since reliable alternatives are available.

Second, we endorse reporter assays not subject to false positives to evaluate potential IRES activity. The Tornado system reliably rejects false-positive IRESes (Unti *et al*, 2024), and we found the HCV IRES produced over 1,000-fold more luciferase than both random sequence negative controls and the toolbox study’s “cellular IRESes” (Fig. 4). Curiously, the *Hoxa9* and *Chrdl1* promoters produced slightly more luciferase than random sequences in the Tornado system. It is possible that this results from rare *trans*-ligation of LgBiT-only transcripts from the *Hoxa9* and *Chrdl1* promoters to SmBiT containing transcripts from the plasmid’s CMV promoter, analogous to *trans*-splicing in the ZKSCAN1 plasmid. Direct transfection of linear mRNA, considered a gold standard for IRES studies (Terenin *et al*, 2017; Loughran *et al*, 2025), is somewhat more difficult but provides unambiguous results. Using A-cap / stemloop linear mRNA transfection, we found three candidate IRESes from the toolbox study had no IRES activity (Fig. 7). Notably, while our work was in review, a preprint was posted that tested several of the same candidate IRESes as *in vitro* transcribed circRNA transfected into HEK293T cells (Koch *et al*, 2026). While *Hoxa9* circRNA that contained linear contaminants appeared to have weak IRES activity (preprint Figure 3C), removal of these contaminants eliminated this activity (preprint Figure 4C). These results are consistent with our findings, in that there was no significant IRES activity for purified circRNA containing the *Hoxa9* promoter. Direct RNA transfection also allows researchers to quantitatively compare m7G-cap dependent translation to IRES activity (Fig. 7). This comparison can help researchers evaluate the potential biological significance of candidate IRES activities. Including appropriate positive (e.g. validated viral IRESes) and negative (e.g. length-matched scrambled sequences without uORFs) controls is crucial. With such controls, the combination of the Tornado plasmid and direct mRNA transfections provide a reliable platform to compare candidate IRESes to known viral IRESes and cap-dependent translation.

Finally, we promote combining m7G-cap dependent sequencing methods (e.g. nAnT-iCAGE) and long read sequencing (e.g. PacBio Iso-Seq) to annotate transcriptomes. In recent years, full-length cDNA sequencing has emerged as the best available technology for identifying mRNA transcripts, especially when paired with orthogonal, high-resolution cap-dependent data like nAnT-iCAGE (Pardo-Palacios *et al*, 2024b). The toolbox study claimed that PacBio Iso-Seq is unreliable due to purported RT drop-off leading to truncated 5’ ends. However, this assertion was based solely on a selective manual comparison of 5’ ends that were shorter than transcript models listed in their chosen GENCODE annotations. We systematically compared their data to nAnT-iCAGE transcription start sites from equivalent mouse embryonic tissue (Ivanov *et al*, 2022) and found that the toolbox study’s PacBio Iso-Seq data accurately quantitated mRNA 5’ ends with single-nucleotide resolution. Our analyses show that PacBio Iso-Seq mRNA annotations can be highly reliable, though they should be cross referenced against orthogonal datasets like nAnT-iCAGE to confirm transcription start sites. We recommend confirming 5’ UTRs using public TSS databases and PacBio Iso-Seq datasets before testing their functions.

While Isoquant can identify novel transcripts, it relies heavily on user-provided annotations (Pardo-Palacios *et al*, 2024b), and so should be used cautiously. The toolbox study provided Isoquant with two incorrect *Hoxa9* annotations, “A” and “B”, that are much longer and much shorter, respectively, than transcripts reported by northern blotting (Rubin *et al*, 1987; Fujimoto *et al*, 1998). We reran Isoquant without *Hoxa9* annotations to allow the program to generate *de novo* transcript predictions. This identified transcripts that fit the published *Hoxa9* mRNA length, accounting for polyA tails (∼250 nt)(Kühn *et al*, 2009). Our results show that providing incorrect transcript annotations to Isoquant creates errors when annotating transcripts with PacBio data. Other programs do not rely as heavily on user provided transcript annotations (Pardo-Palacios *et al*, 2024b) and may thus be more reliable. For example, the SQUANTI3 package (Pardo-Palacios *et al*, 2024a) uses TSS peak locations defined by CAGE data to refine and validate PacBio Iso-Seq transcript models and was recently incorporated into PacBio’s recommended workflow.

Our Isoquant analysis also detected a pri-mir196b related non-coding RNA expected to be retained in the nucleus, just as we previously proposed based on similar data (Akirtava et al., 2022). Similar cleavage products of other pri-miRNA transcripts are commonly observed in EST sequences (Krzyzanowski *et al*, 2011). We show pri-mir196b is targeted by the toolbox study’s “IRES” smFISH probe, which can explain why this smFISH probe signal is only found in nuclei, in the same embryonic regions where mir196 miRNAs are expressed (Wong *et al*, 2015). While smFISH is useful for detecting the presence and location of RNA, it detects both mRNA and non-coding RNA. When non-coding RNAs and mRNAs share sequences, as with *Hoxa9* mRNA and pri-mir196b lncRNA, this lack of specificity makes it impossible to discern which RNA was detected by smFISH probes. In contrast, methods that are highly selective for mRNA, e.g. PacBio Iso-Seq, provide reliable detection and annotation of mRNAs. Despite its specificity for m7G-capped polyadenylated transcripts, PacBio Iso-Seq captured the pri-miR196b ncRNA. Close inspection reveals that the Iso-Seq reads supporting pri-mir196b terminate in poly-A stretches encoded in the genome that likely allow internal initiation of reverse transcription. A small fraction of pri-mir196b transcripts may be m7G-capped after processing, which would allow full length cDNA generation. Reads corresponding to unspliced *Hoxa9* mRNA also terminate in the same internal genomic polyA stretch, suggesting these reads represent pre-mRNAs. Thus, it should be noted that PacBio Iso-seq can amplify non-coding RNAs and pre-mRNAs that contain internal poly-A segments.

Although hundreds of cellular IRESes have been reported over three decades using plasmid assays, direct mRNA transfection experiments routinely fail to validate them (Terenin *et al*, 2017; Yang & Wang, 2019). A common explanation for this is that cellular IRESes require a “nuclear experience”. However, the required “nuclear experience” is likely to be the creation of artifacts by promoters and splice sites. The Tornado plasmid may provide clarity, as it expresses luciferase through a “nuclear experience” yet has much lower background activity from promoters and does not involve spliceosomal splicing. Critically, 5’ UTR annotations often include active, physiologically relevant promoters and splice sites, due to both annotation errors and the complex architecture of mammalian genes (Akirtava *et al*, 2022), making them difficult to define. The approaches promoted by the toolbox study – smFISH and qRT-PCR, which are not specific for mRNA, and a back-splicing circRNA plasmid susceptible to promoter artifacts – exacerbate these problems. Here, we show: (1) full length cDNA sequencing, backed by orthogonal nAnT-iCAGE data, provides accurate mRNA annotations and (2) that direct mRNA transfection and the Tornado circRNA plasmid provide accurate tests of IRES activity. Used with care, these approaches will allow future IRES researchers to move forward with confidence.

## Methods

### mRuby3-ZKSCAN-eGFP reconstruction and cloning

The sequence of the mRuby3-ZKSCAN-eGFP plasmid (Chen *et al*, 2021) was provided by Dr. C.K. Chen. The plasmid was recreated by modification of the pcDNA3.1(+) ZKSCAN1 Sense plasmid (Addgene #60631). A fragment containing the downstream Alu element was PCR amplified using the primers Vect-F3-PstI-F and Vect-frag-3-R (Table S2) and digested with PstI. A gene fragment containing the CMV promoter, mRuby, the upstream Alu element, and split GFP was synthesized (Twist Biosciences) and digested with PstI. The two fragments were ligated, and the ligated product was PCR amplified using the primers TWLF-F and TWLF-R. The subsequent product was digested with BglII and NotI and cloned into pcDNA3.1(+) ZKSCAN1 to generate the mRuby-ZKSCAN1-split-eGFP reporter construct. Mouse *HOXA9* and *CHRDL1* promoters were amplified from previously cloned vectors (Akirtava *et al*, 2022) using the primers MMHoxa9-Zkscan-F/R, and MMChrdl1-Zkscan F/R respectively (Table S2). The PCR products and mRuby-ZKSCAN1-split-eGFP reporter construct were digested with EcoRV and the HOXA9 and CHRDL1 promoters were cloned into the plasmid using In-Fusion® Snap Assembly Master Mix (Takara) to generate the vectors mRuby-ZKSCAN1-split-eGFP-HOXA9, and mRuby-ZKSCAN1-split-eGFP-CHRDL1. Vector sequences were verified by full-plasmid sequencing.

### mRuby3-ZK-spEGFP trans-splicing assays

A 6 well plate was seeded with 2 x 10^5^ HEK293T cells per well in 2 ml of DMEM supplemented with 10% FBS. The cells were and incubated overnight at 37°C. Each well was transfected with 2.5□g mRuby-ZKSCAN1-split-eGFP-HOXA9, mRuby-ZKSCAN1-split-eGFP-CHRDL1, or a 1:1 mixture of both reporters (1.25□g each), using Lipofectamine™ 3000 (Invitrogen). The cells were incubated for 24 hours and harvested by removing the media and adding 1 ml of TRIzol™ (Invitrogen™) to each well. Total RNA was extracted according to the manufacturer’s instructions and precipitated with 1 volume of isopropanol. Resuspended RNA (225 l of nuclease free water) was treated with 2 units of TURBO DNase™(Invitrogen™) for 1 hour at 37°C. The RNA was re-extracted with 500□l of acid phenol chloroform, precipitated with 1 volume of isopropanol, and resuspended in 100 l of nuclease free water. For the co-transfection experiment, 5□g of total RNA was poly A selected using the NEBNext® Poly(A) mRNA Magnetic Isolation Module (New England Biolabs) according to the manufacturer’s instructions in Fig. 1E. To digest linear (but not circRNA), 2.5□g of total RNA and 50 ng of poly A selected RNA was digested with 2.5 units and 0.05 units of RNase R (New England Biolabs) respectively for 15 minutes at 37°C. The reaction was stopped by adding EDTA to 10 mM and the RNA was purified over RNA Clean & Concentrator -5 column (Zymo Research). Both RNase R treated and untreated RNAs were subjected to RT-PCR using the OneTaq® RT-PCR kit (New England Biolabs) according to the manufacturer’s instructions (for primer sequences see Table 1). To control for possible strand switching during RT-PCR, RNA from the *HOXA9* and *CHRDL1* transfections were mixed 1:1, as a template for RT-PCR reactions detecting *HOXA9* and *CHRDL1 trans*-spiced fusion products. Prior to PCR, the cDNA were purified using SPRI magnetic beads and resuspended in 50 l of nuclease free water. For each reaction, 2 l cDNA was amplified for 30 cycles with a 53°C annealing temperature. Amplification was assessed using 1 l of PCR product on an automated electrophoresis TapeStation D1000 tape (Agilent).

### RNA-seq from plasmid transfected cells

HEK293T cells (5 x 10^4^) were seeded in a 6-well dish in DMEM with 10% FBS. The cells were incubated overnight, and transfected with 500 ng of the Back-Splicing and Tornado plasmids carrying candidate *Hoxa9, Chrdl1*, and *Dlx1* IRESes in duplicate, using 8 μl Lipofectamine™ 3000 (Invitrogen) according to the manufacturer’s instructions. After 24 hours, total RNA was extracted by using 500 μl TRIzol™ (Invitrogen™). The RNA was purified using an RNA Clean & Concentrator column (Zymo Research) and eluted in 50 μl of nuclease free water. The RNA integrity was confirmed using automated gel electrophoresis, and 1 μg of total RNA was sent to an external facility for polyA+ RNA sequencing library preparation (NEB Ultra II) and sequencing. In addition, 1 μg of total RNA for one replicate was sent for rRNA depletion sequencing (Qiaseq FastSelect -rRNA HMR kit). The resulting sequencing reads were aligned to the human genome (hg38) with transcriptome annotations (MANE) using hisat2 with default options. Reads aligned to MANE annotated transcripts were identified and counted using bedtools coverageBed (Quinlan & Hall, 2010). Reads mapping to splice junctions from the back-splice plasmid were identified and counted using regtools junction extract with default parameters (Cotto *et al*, 2023). RPKM values were calculated for all MANE transcripts and plasmid splice-junctions by dividing the mapped reads by the length of the target transcript or splice junction, divided by the total number of mapped reads (in millions) per sample.

To verify the presence of Tornado plasmids circRNA junction sites, 500 – 900 ng of total RNA was reverse transcribed in a 20 μl reaction for 1 hour at 42 °C with a reverse primer that anneals to small bit nano luciferase (sNLUC-R) using M-MuLV reverse transcriptase. The resulting cDNA was bead-purified and resuspended in 50 μl of 1x TE. 2 μl of cDNA was amplified in a 20 μl reaction for 30 cycles using OneTaq® Hot Start PCR mix, with the primers HOXA9-UTR-F/sNLUC-R, DLX1-UTR-F/sNLUC-R, CHRDL1-UTR-F/sNLUC-R, using a 55 °C annealing temperature. The PCR products were run on a 1% agarose gel, excised, and sent for Sanger sequencing with both the forward and reverse primers for each product.

### Luciferase vector construction for RNA transfection

A modified version (BglII restriction sites deleted) of the phMGFP plasmid (Promega) was used as a vector backbone. A T7 promoter followed by a hairpin was synthesized for A-cap / stemloop IRES constructs and PCR amplified with the primers T7-Stem-insert-F and T7-Stem-insert-R (Table S2). The resulting PCR product was digested with SacI and HindIII and cloned into the phMGFP vector upstream of GFP, resulting in the vector T7HP-GFP. Separately, a T7 promoter lacking the stemloop was generated. Overlapping oligos T7-Gcap-F and T7-Gcap-R were phosphorylated, annealed, and cloned into phMGFP digested with SacI and HindIII, resulting in the vector T7-GFP. Both plasmids were sequence verified. GFP was removed from T7-GFP and T7HP-GFP by digestion with BglII and NotI, and replaced with nanoluciferase (n-luc) amplified from the pmirGLO vector (Promega) using primers N-luc-BglII-F and N-luc-NotI-R, resulting in the vectors T7-nluc and T7HP-nluc. The EMCV, HCV, and CrPV IRES sequences were amplified from corresponding plasmid reporters (Russell *et al*, 2023) using PCR primers (Table S2) and cloned into the T7-nluc and T7HP-nluc vectors digested with HindIII and BglIII. EMCV was cloned using the In-Fusion® Snap Assembly Master Mix (Takara), while HCV and CrPV were cloned by HindIII / BglII digestion and ligation. The mouse *Chrdl1* and *Hoxa9* promoter (putative IRES) sequences were amplified from plasmids (Akirtava *et al*, 2022) using primers flanked by HindIII and BglII sites, and cloned by ligation. The shorter Hoxa9 UTR (85 nt), was amplified from the *Hoxa9* plasmid with primers MM-Hoxa9-real-HindIII-F and N-luc-BglII-R. The *CDK11a*, *Dlxl*, and three random sequences generated by scrambling the *CDK11a* sequence were synthesized as gene fragments (Twist Biosciences) containing 5’ HindIII and 3’ BglIII sites. After PCR amplification, inserts were digested with HindIII and BgllII and cloned into T7-nluc and T7HP-nluc. All constructs were sequence verified by full-plasmid sequencing (sequences available in Extended Data).

### Tornado-split-nLuc vector cloning

To clone UTR sequences upstream of a split nano luciferase, the pcDNA3.1+-Tornado-split-nLuc vector (Addgene #212612 (Unti *et al*, 2024)) was digested with EcoRI and BsiWI. The inserts were PCR amplified from corresponding T7-nluc constructs with primers that added EcoRI and BsiWI sites. The PCR products were digested with EcoRI and BsiWI and cloned into the vector described above.

### Tornado-split-nLuc Luciferase assays

HEK293T cells and mouse C3H/10T1/2 cells were plated at 5 x 10^3^, and 2.0 x 10^3^ cells respectively, in 100 l of DMEM / 10% FBS in each well of a 96 well plate and incubated overnight at 37°C. HEK293T cells were transfected for 24 hours with 75 ng of each reporter construct per well with 45 ng of a control vector expressing Firefly luciferase from the mouse *Hoxa5* promoter (Akirtava *et al*, 2022), using Lipofectamine™ 3000 transfection reagent (Invitrogen). C3H/10T1/2 cells were transfected for 24 hours with 75 ng of each reporter construct per well with 45 ng of a control vector expressing F-luciferase. Due to the toxicity of lipofectamine to C3H/10T1/2 cells, the cells were instead transfected with ViaFect™ Transfection reagent (Promega). Six replicates were performed for each construct. The Nano-Glo® Dual-Luciferase® Reporter Assay System (Promega) was used to measure N-luc and F-luc luminescence, according to the manufacturer’s instructions, on a Tecan microplate reader. Data were processed by subtraction of luminescence values from untransfected negative control wells and N-luc / F-luc ratios were calculated for each replicate. Replicate data are shown in Table S3.

### A cap / stemloop and m7G cap mRNA Luciferase assays

Luciferase reporter constructs were PCR amplified with the primers phMGFP-T7-F and phMGFP-GFP3p-R to generate T7 transcription templates. 250 ng of each template was used for RNA synthesis, using the HiScribe® T7 Quick High Yield RNA Synthesis kit (New England Biolabs) according to the manufacturer’s instructions. The ARCA m^7^G-cap analog (New England Biolabs; S1411) was used at a 4:1 ratio to GFP for transcription of T7-nluc constructs, while an A-cap analog (New England Biolabs S1406), was used to generate T7HP-nluc IRES reporter RNAs. RNAs were ethanol precipitated and resuspended in 100 l of nuclease free water. The RNA integrity was verified by gel electrophoresis on an Agilent Tapestation and quantified using a Qubit™ RNA high sensitivity assay (Invitrogen). HEK293T cells and mouse C3H/10T1/2 embryonic mesenchymal cells were plated at 1 x 10^4^, and 4.0 x 10^3^ cells respectively, in 100 l of DMEM / 10% FBS in each well of a 96 well plate. The cells were incubated overnight at 37°C and transformed at 50-70% confluency. For each well, 30 ng of each N-luc RNA was co-transfected with 10 ng of Firefly luciferase (F-luc) control RNA using Lipofectamine™ MessengerMAX ™ Transfection Reagent (Invitrogen) for four hours. Four technical replicates were performed for each construct. The Nano-Glo® Dual-Luciferase® Reporter Assay System (Promega) was used to measure N-luc and F-luc luminescence, according to the manufacturer’s instructions, on a Tecan microplate reader and the results were quantified as described above. Replicate data are available in Table S3.

### PacBio Iso-Seq and nAnT-iCAGE data processing and 5’ end correlation analyses

nAnTi-CAGE reads (Ivanov *et al*, 2022) were processed and mapped to the mm39 genome as previously described (Akirtava *et al*, 2022). Full length PacBio Iso-Seq reads were selected by matching both the 5’ template switching oligo and the oligo-dT 3’ RT primer sequence. This is equivalent to processing using the “lima” module recommended by PacBio (https://github.com/PacificBiosciences/IsoSeq). Raw reads were first filtered to select 5’ TSO containing reads and remove the TSO sequence (CAATGAAGTCGCAGGGTTGGG; “PBF” in Fig. 1) using cutadapt ((Martin, 2011) version 4.5; options -j 12 --trimmed-only -a CCCAACCCTGCGACTTCATTG -g CAATGAAGTCGCAGGGTTGGG -O 15). A custom perl script (getIso-SeqStrandFL.pl) was then used to identify the polyA sites on TSO-trimmed reads. Reads containing 5’ polyA sites were reverse complemented to provide their sense orientation. The resulting trimmed reads were then mapped to the mm39 genome using minimap2 ((Li, 2018) version 2.17-r941; options -ax splice:hq -u f -t 12). Deeptools bamCoverage ((Ramírez *et al*, 2016) version 3.51; options --exactScaling binSize 1 --Offset 1 1 --filterRNAstrand = (forward/reverse)) was used to produce strand-specific bigwig files mapping 5’ ends of PacBio Iso-Seq alignments and of nAnTi-CAGE reads. Bigwig files were visualized at individual loci using the Integrated Genome Viewer (Robinson *et al*, 2011) and browser regions were exported as svg files to generate Fig. 6.

To quantitate correlations between PacBio and nAnTi-CAGE read counts around transcript 5’ ends, bigwig files were converted to bedGraphs using the bigwigTobedGraph tool (Kuhn *et al*, 2013). Regions 500 nts up and downstream of annotated transcript 5’ ends (NCBI Refseq, mm39, March 15, 2025) were intersected with nAnTi-CAGE and PacBio 5’ ends using intersectBed (v2.30.0 (Quinlan & Hall, 2010)). The resulting files were processed using a custom Perl script (1kbTSSregioncollapsePearson.pl) to calculate read numbers and Pearson’s R correlation coefficients for 5’ ends using individual nts and, separately, using a 5-nt sliding window (Table S2). Transcripts were filtered to remove low expressed transcripts (RPKM < 1). The plots in Fig. 5 used Pearson’s R values with 1-nt resolution comparisons and were generated using R (version 4.2.0).

### Isoquant analysis without user-provided *Hoxa9* and *Hoxa10* transcript annotations

PacBio Iso-Seq data from Koch et al. were aligned to the mouse genome (mm39) using Isoquant (version 3.6.3; (Prjibelski *et al*, 2023) with options --complete_genedb --data_type pacbio_ccs options). A custom genome annotation file lacking the *Hoxa9* and *Hoxa10* annotations from GENCODE version M36 (Ensembl 113) was used as input for Isoquant mapping. Isoquant aligned reads using minimap2 (version 2.28-r1209; (Li, 2018)) with the raw PacBio Iso-Seq reads as input. The resulting Isoquant transcriptome annotation file predicted a novel gene (“novel_gene_chr6_3912”) at the *Hoxa9* locus containing the five transcripts depicted in Fig. 6. These novel transcripts were visualized in IGV, along with the minimap2 alignment of full-length trimmed PacBio Reads (generated in the correlation analysis above). Reads were manually sorted into groups corresponding to each Isoquant isoform and counted to estimate their relative expression level (as shown in Fig. 6).

### Image processing

Koch et al.’s smFISH image files from Fig. 3D, “D1.tif”, “D2.tif”, “D3.tif”, “D5.tif”, “D6.tif”, and “D7.tif” were downloaded from their data host website. Images were processed in Fiji (version 2.16.0) (Schindelin *et al*, 2012) to produce the composite images in Fig. 3 by merging the blue (DAPI), red (“IRES” probe) and green (“CDS” probe) channels. smFISH puncta were identified simultaneously for red and green probes using the ComDet plugin (v 0.5.6) (Katrukha *et al*, 2021) with a maximum distance between ovals of 2 pixels, minimum size of 3 pixels, and minimum intensity of 20 SD. Puncta inside the nucleus (determined by thresholding the blue channel of DAPI images from 35 to 255), were calculated separately. The numbers of nuclear and cytoplasmic spots were calculated and plotted in Fig. 7.

### Polysome profiling and qRT-PCR

A 10 cm dish was seeded with 1 x 10^6^ HEK293T cells in 10 ml of DMEM with 10% FBS and incubated overnight at 37°C. Cells were harvested by removing the media and resuspended in 10 ml of ice-cold PBS with 100 g/ml cycloheximide using a cell scraper. The cell suspensions were transferred to 15 ml conical tubes and pelleted for 1 minute at 500 G. The supernatant was decanted, and the cells were resuspended in 1 ml of PLB (50 mM Tris-HCL pH 7.5, 140 mM KCl, 12 mM MgCl_2_, 1% Triton-X 100, 1 mM DTT, cOmplete™ EDTA-free protease inhibitor cocktail (Roche), 500□g/ml cycloheximide, 0.2 U/□l SUPERase-In™ (Invitrogen), and 0.1 U/□l

RNasin® Ribonuclease inhibitor (Promega). The cells were allowed to lyse on ice for 10 minutes and then transferred to 1.5 ml tubes. The lysate was centrifuged at 20,000 G for 10 minutes at 4°C and the supernatant was removed to a new tube. 100 ng of *in vitro* transcribed scramble-1 nano luciferase RNA was added to the lysate, briefly mixed, and 400□l of lysate was centrifuged over a 10 – 50% sucrose gradient (in PLB lacking Triton-X 100) and fractionated into 14 fractions using a Biocomp polysome fractionator.

Total RNA was extracted from 200 l of each fraction (and unfractionated lysate) using 500□l TRIzol™ (Invitrogen™), purified using an RNA Clean & Concentrator column (Zymo Research) and eluted in 50□l of nuclease free water. cDNAs were synthesized in 20□l reactions at 55°C for 30 minutes from 5□l total RNA with the primers N-luc-seq-R and HSGAPDH-R using Superscript IV (Invitrogen). The cDNAs were purified with 1.0X PCR purification beads and eluted in 20□l of nuclease free water. Equal volumes of cDNAs were combined from corresponding fractions to represent top, 40S, 60-80S, light (2-5 ribosomes), medium (6-9 ribosomes) and heavy (9+), and nuclease free water was added to equalize the volumes of the fractions respective to each other. Using the primers SCR1 F/R or HSGAPDH F/R, 2□l of cDNA was amplified using the DyNAmo Flash SYBR Green qPCR kit (Thermo Scientific™) for 40 cycles with an annealing/extension/capture step at 60°C for 15 seconds. Four technical replicates were performed for each cDNA and amplicon as well as four replicates of no template controls. The relative abundance of RNA in each fraction was estimated using the□CT method (Panda *et al*, 2017). RNA abundance was estimated as 2^(-CT_fraction_ – max(□CT_replicate_)). For each technical replicate, this approach sets the expression value of the fraction with the lowest RNA expression to 2^0 = 1, with all other fractions expressed as a multiple of the minimum value. The expression values for each fraction were then calculated as a percentage of the total expression values for fractions from each replicate.

## Extended data

1. File S1 - Plasmid vector sequences

2. Table S1 – RNA-seq data from plasmid transfections

3. Table S2 – Primers and plasmid insert sequences

4. Table S3 – Luciferase assay data.

5. Table S4 – nAnT-iCAGE vs PacBio Iso-seq TSS data

## Data Availability

Raw RNA-seq data have been deposited at NCBI under BioProject ID: PRJNA1526329 and will be released upon publication.

## Conflict of interest statement

The authors declare no conflict of interests.

## Supporting information

Plasmid and insert sequences

Isoquant de novo transcript models.

New Table S1 (relating to figure 2)

Now Table S2

Now Table S3

Now Table S4

## Acknowlegements

The authors would like to thank members of the McManus lab and Pasha Baranov, Angela Brooks, Tom Dever, Sergey Dimitriev, Rachel Green, Alan Jacobson, Samie Jaffrey, Craig Kaplan, Gary Loughran, Andrey Prjibelski, Leos Valasek and Jeremy Wilusz for helpful conversations, advice and suggestions related to this work. We thank Howard Chang and C.K. Chen for providing the full sequence of the ZKSCAN1 back-splicing reporter plasmid from Chen et al., 2021. This work was supported by NIH grant R35GM145317 to CJM.

## Supplemental Figures

**Figure EV1.**
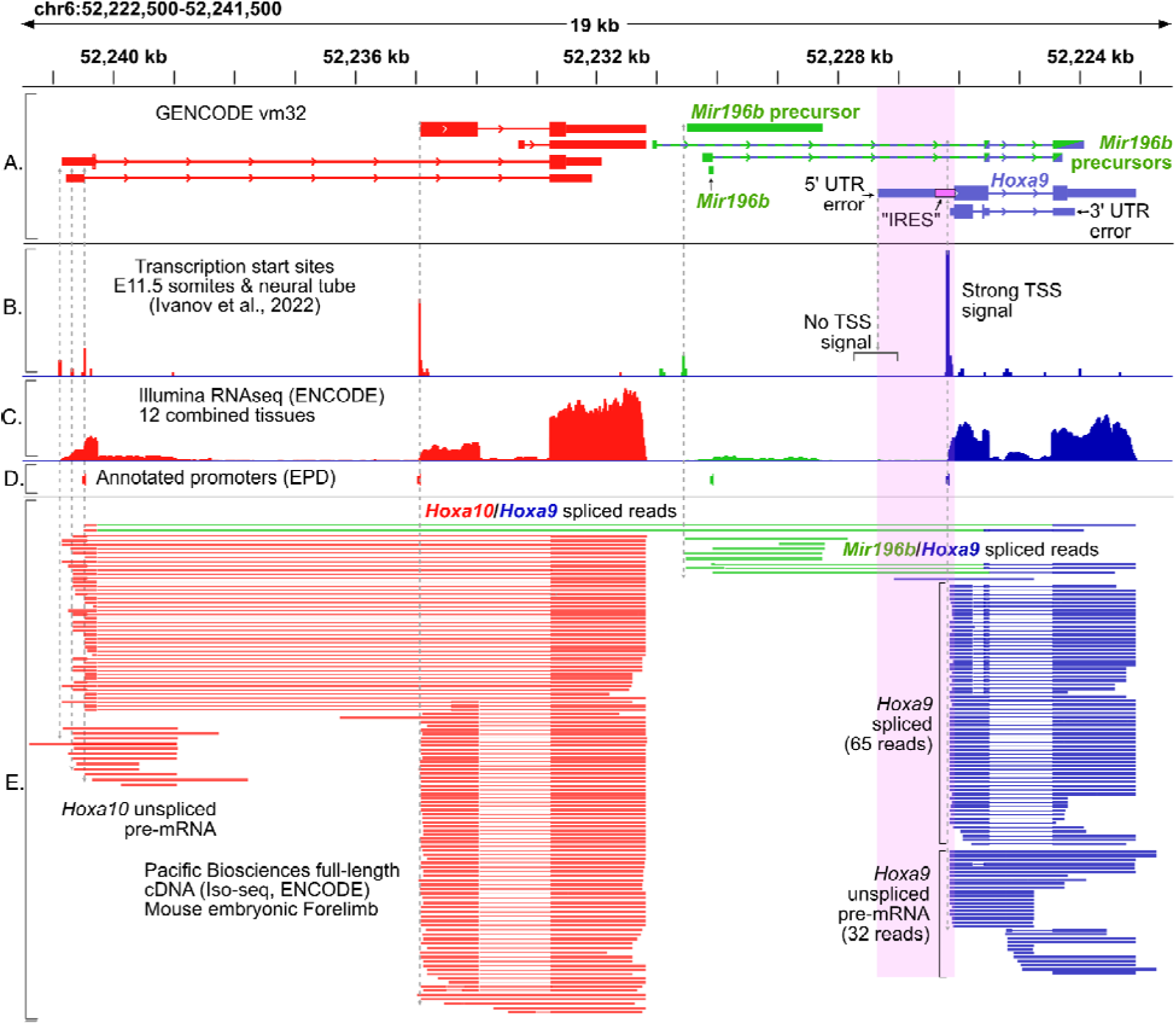
RNA-seq from mouse embryos shows transcription from Hoxa10 and miR196b extending through the Hoxa9 promoter and coding region. Genome browser tracks showing the Hoxa10, Mir196b, and Hoxa9 loci in the mouse genome (mm10). The genomic coordinates have been oriented to show the region from 5’ to 3’. **(A)** GENCODE (version m32) transcript annotations are shown, including Hoxa10 (red), Mir196b (green), precursor transcripts for mir196b spliced to Hoxa9 (green / blue), and Hoxa9 (blue). The two Hoxa9 isoforms are misannotated due to errors in their 5’ and 3’ UTR coordinates. **(B)** nAnTi-CAGE sequencing data mapping m7G-capped 5’ transcription start sites (TSS) are plotted (Ivanov et al., 2022), showing the major transcription start sites (dashed gray arrows). Note the total absence of transcription start site data at the misannotated “IRES” isoform of Hoxa9. Illumina **(C)** RNA-seq data combined from twelve mouse tissues (Akirtava et al. 2022). The RNA-seq signal increases immediately downstream of the somite and neural tube Hoxa9 transcription start site, with scant mRNA signal upstream in the 5’ UTR of the “IRES” isoform. **(D)** Eukaryotic promoter database promoter annotations. **(E)** PacBio Iso-Seq full-length cDNA reads from mouse embryonic forelimb are shown. Two reads show Hoxa10 - Hoxa9 fusion transcripts generated by intron splicing. Multiple reads correspond to Mir196b precursor transcripts (green), including Mir196b transcripts spliced to Hoxa9 sequences (green / blue). The misannotated Hoxa9 5’ UTR and promoter region overlaps pri-mir196b precursor transcripts and introns from Hoxa10/Hoxa9 fusion transcripts and precursor transcripts from Mir196b (pink rectangle). Adapted from Akirtava et al., 2022.

**Figure EV2.**
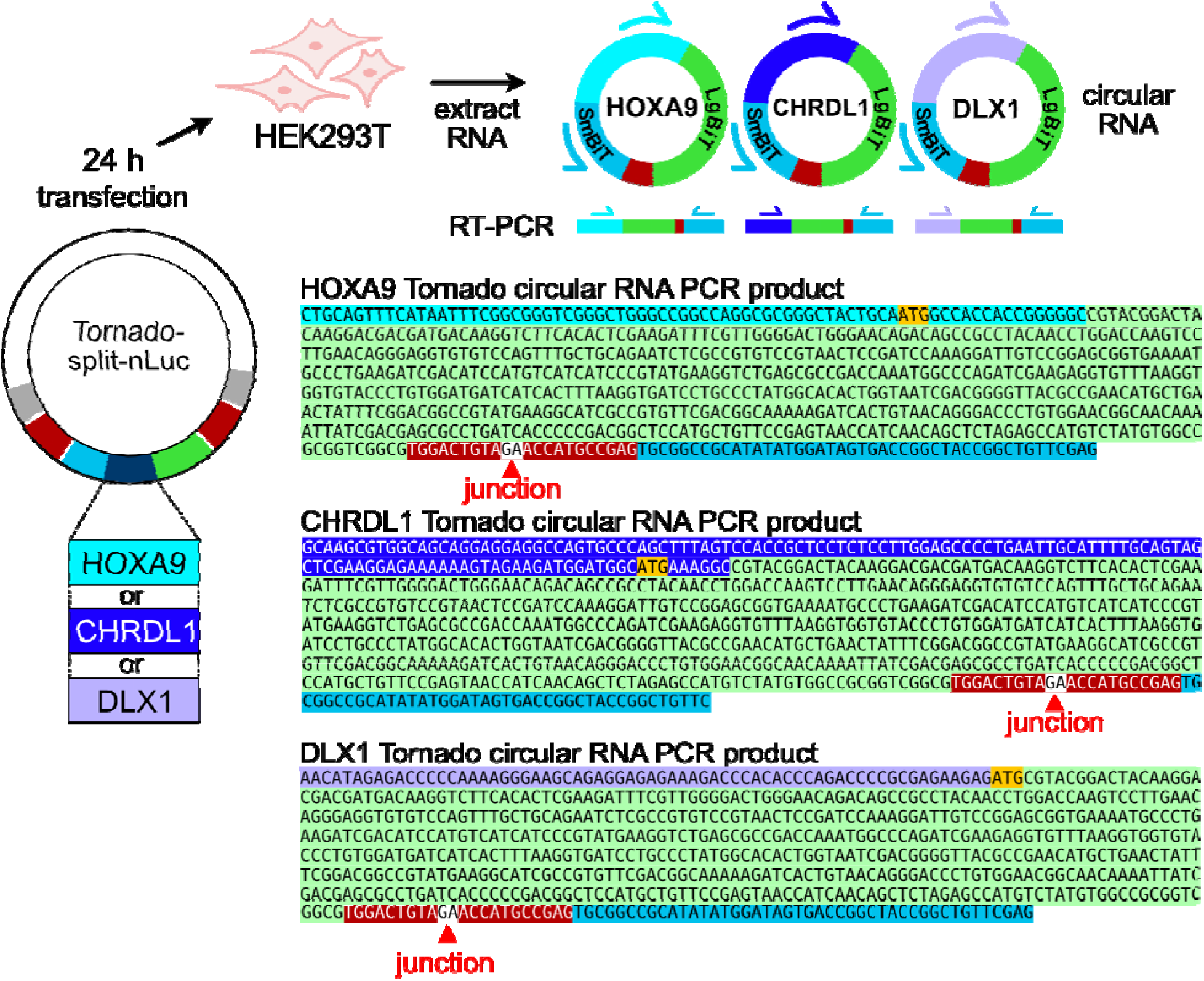
The Tornado split n-luc plasmids produce transcripts with correct luciferase ligation junctions. Plasmids with the HOXA9, CHRDL1, and DLX1 promotors cloned into the IRES test region were transfected for 24 hours into HEK293T cells. Total RNA was extracted and the circular RNA products were reverse transcribed with a primer that anneals to SmBIT. The cDNA was amplified with gene specific forward primers and the SmBIT reverse primer, and the PCR products were sequenced by Sanger sequencing. The resulting sequences are shown above, with the ligation junctions marked with red triangles.

**Figure EV3.**
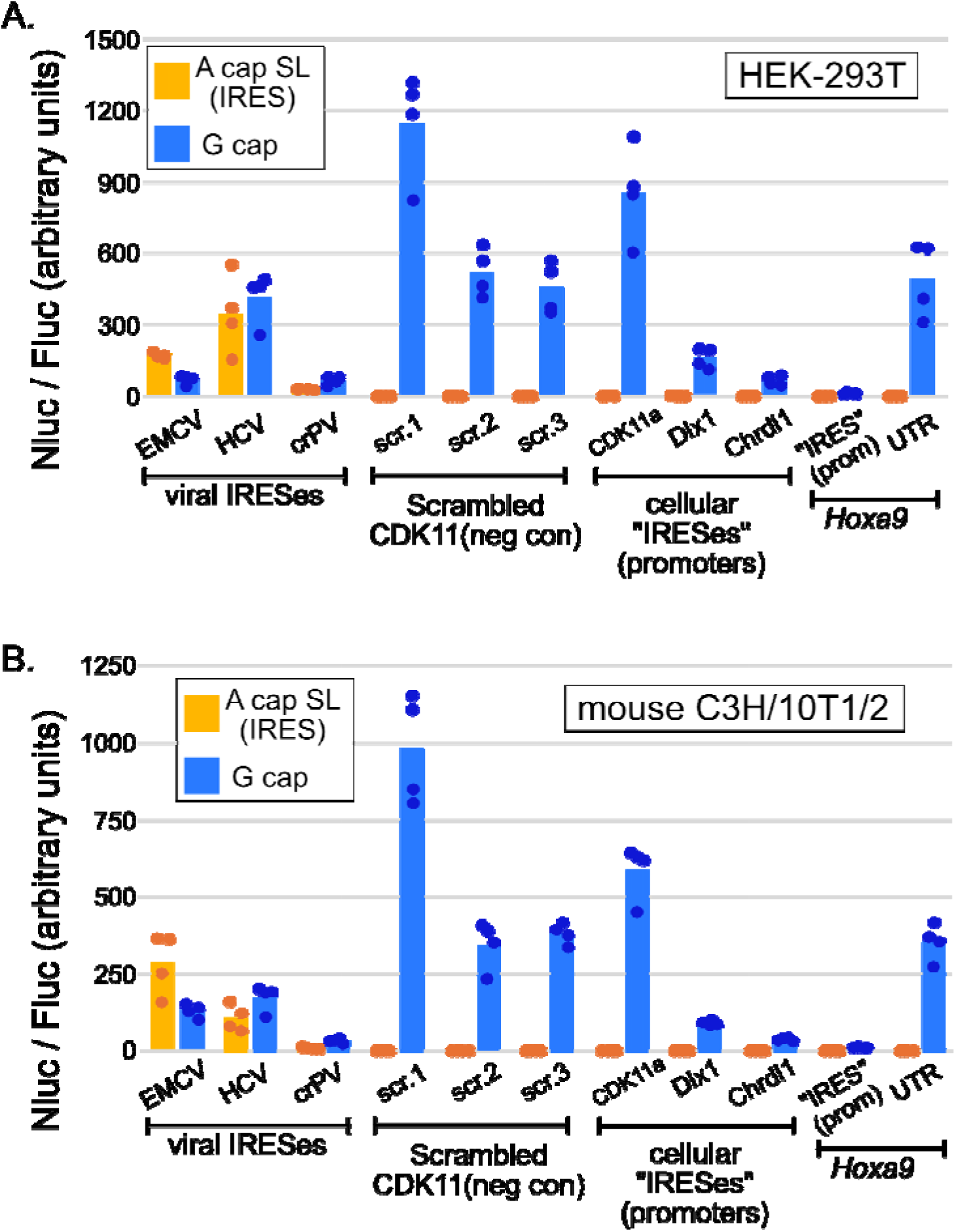
Expanded view of neglligible putative IRES activities in mRNA transfection assays (Main. Figure 4**). A.** Viral IRESes expressed similar levels of Nluc in IRES and G-cap reporters in HEK293T cells. In contrast, the putative cellular IRESes all exhibit IRES activity equivalent to scrambled negative control sequences. The genuine 85 nucleotide Hoxa9 5’ UTR, which lacks all previously reported “IRES” essential sequences, has slightly more IRES activity than the Hoxa9 promoter region claimed to be an “IRES” by the toolbox study G-cap transcripts with the genuine Hoxa9 5’ UTR produced >1000-fold more luciferase than the Hoxa9 “IRES”. Similarly, G-cap Chrdl1 and Dlx1 reporters produce ∼100-fold and ∼200-fold more luciferase than corresponding “IRES” reporters. These plots show the raw data as points on a linear scale (n=4; * = p < 0.05; ** = p < 0.01; Also see Table S3). **B.** Identical experiments from C3H/10T1/2 cells.

**Figure EV4.**
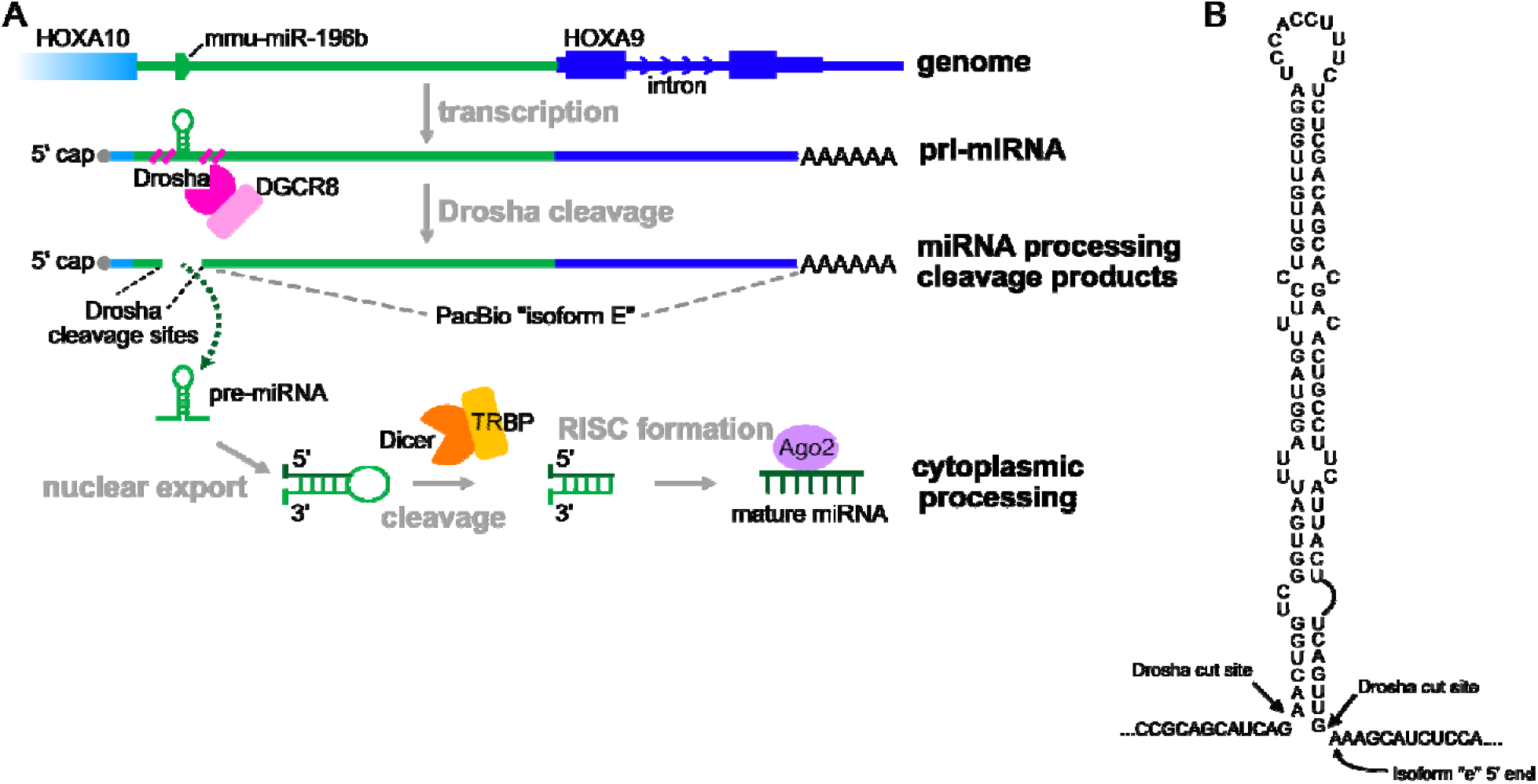
The Isoquant isoform “e” is consistent with canonical processing of a mir196b primary transcript. **A.** Depiction of mir196b processing from fusion transcripts generated from Hoxa10 and/or mir196b promoters that read through the HoxA9 mRNA sequence. The annotated mir196b gene is the precursor RNA, which is removed by Drosha / DGCR8 cleavage. The 5’ end of the resulting trailing product (termed “L2” product) of Drosha cleavage maps precisely to the 5’ end of isoform E identified in PacBio sequence reads using Isoquant de novo. **B.** Zoomed in view of the mir196b primary transcript showing the Drosha cut sites and the 5’ end of Isoquant isoform “e”.

**Figure EV5.**
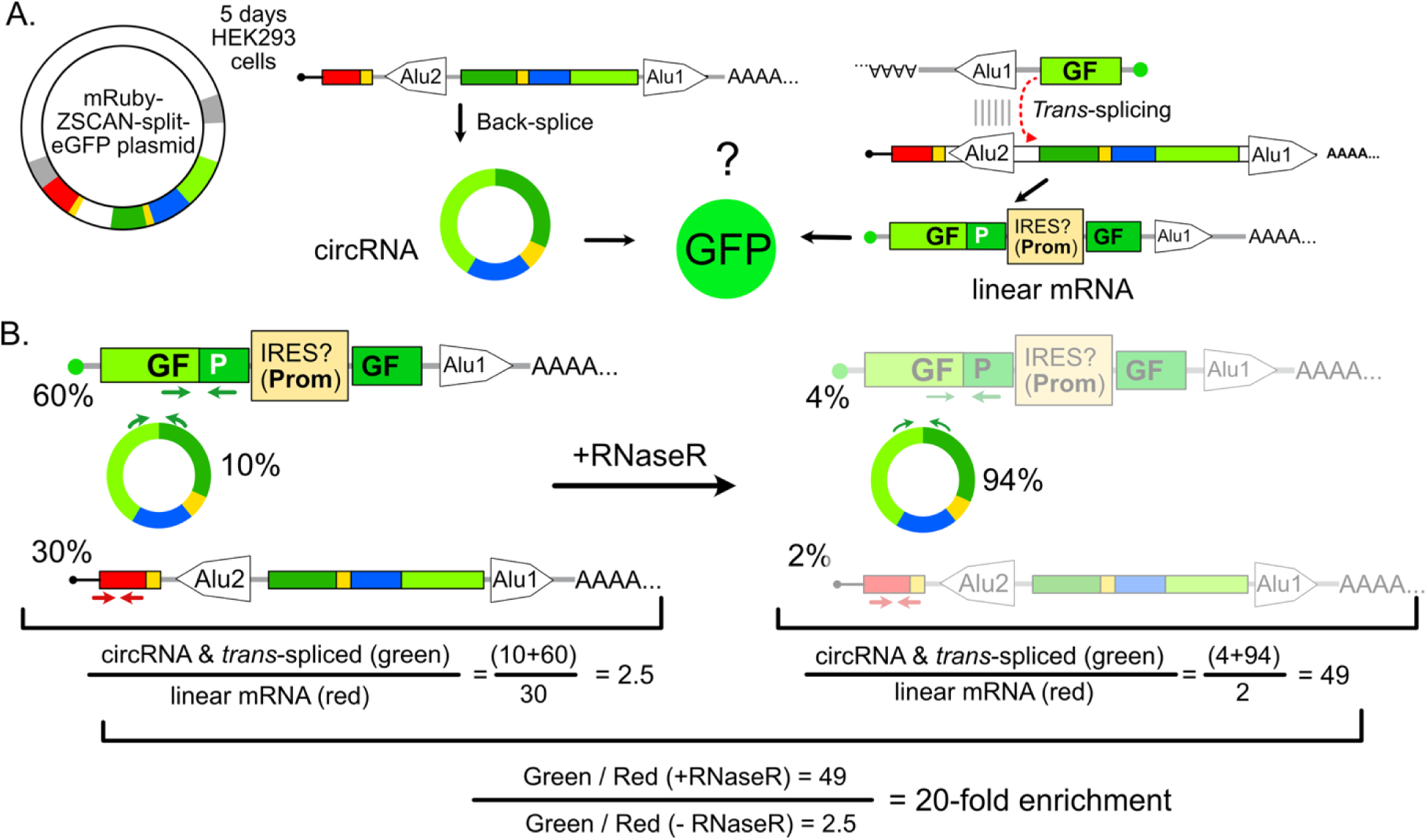
Enrichment of spliced GFP signal vs RFP after RNase R treatment does not rule out the presence of trans-spliced linear GFP mRNA. **(A)** Diagram depicts the circular RNA products produced by back-splicing (left) and the monocistronic linear GFP mRNA artifacts products produced by trans-splicing (right). CircRNA can produce GFP if the insert sequence (blue) has IRES activity. The linear mRNA produced by trans-splicing could produce GFP by cap-dependent scanning. If there are any trans-spliced artifacts, it is impossible to know whether GFP is expressed due to IRES activity or due to cap-dependent scanning. **(B)** The qRT-PCR assay that Koch et al. used in attempts to show GFP is expressed from circRNAs used primers across the GFP splice junction (green) and in mRuby (red). GFP primers amplify both trans-spliced contaminants and circRNA, while mRuby primers amplify only the linear pre-mRNA. RNase R treatment selectively depletes both the linear mRuby and the linear trans-spliced monocistronic GFP mRNAs. The result is an enrichment of signal from the GFP amplicon, even when trans-spliced linear contaminants are more abundant than circRNA. The assay only indicates that circular RNAs were present. It does not rule out the presence of trans-spliced contaminant artifacts.

## Notes

### Competing Interest Statement

The authors have declared no competing interest.

### Summary of Updates

A new figure (2) was added to quantify the levels of spliced linear polyAdenylated GFP expressed from the ZKSCAN1 back-splicing reporter plasmids. Additional supplemental figures were added to clarify other aspects of the manuscript, and additional references were cited and discussed.

